# Iron deficiency causes aspartate-sensitive metabolic and proliferative dysfunction in CD8+ T cells

**DOI:** 10.1101/2024.02.01.578381

**Authors:** Megan R. Teh, Nancy Gudgeon, Joe N. Frost, Linda V. Sinclair, Alastair L. Smith, Christopher L. Millington, Barbara Kronsteiner, Jennie Roberts, Bryan P. Marzullo, Alexandra E. Preston, Jan Rehwinkel, Thomas A. Milne, Daniel A. Tennant, Susanna J. Dunachie, Andrew E. Armitage, Sarah Dimeloe, Hal Drakesmith

## Abstract

Iron is an irreplaceable co-factor for metabolism. Iron deficiency affects >1 billion people, causing symptoms including anaemia and impaired immunity. Nevertheless, precisely how iron deprivation impacts immune cell function remains poorly characterised. We therefore interrogated how physiologically low iron availability affected activated CD8+ T cell metabolism and function, using multi-omic and metabolic labelling approaches. Iron limitation profoundly stalled proliferation without influencing cell viability, altered histone methylation status and disrupted mitochondrial membrane potential. Consistently, metabolism of glucose and glutamine in the TCA cycle was limited, indeed TCA cycle activity was partially reversed to a reductive trajectory. Previous studies have shown mitochondria-derived aspartate is crucial for proliferation of transformed cells. Surprisingly, we found aspartate was increased in stalled iron deficient CD8+ T-cells, but was not utilised cytosolically for nucleotide synthesis, likely due to trapping within depolarised mitochondria. Conversely, exogenous aspartate, which directly accesses the cytosol, markedly rescued the clonal expansion of even severely iron-deficient CD8+ T-cells. Overall, iron scarcity creates a mitochondrial-located metabolic bottleneck impairing T-cells, which can be bypassed by resupplying inhibited biochemical processes with aspartate. These findings reveal molecular consequences of iron deficiency for CD8+ T cell function, providing mechanistic insight into the basis for immune impairment during iron deficiency.

## Introduction

Studies of metabolism in cancer and in different cell lineages have changed understanding of how metabolic processes regulate cellular transformation and cell fate^1,2^. In contrast, how common nutritional deficiencies influence metabolic pathways and alter cell biology is relatively poorly studied. Iron deficiency is the most common micronutrient deficiency worldwide^3^, inhibiting erythropoiesis, impairing cognitive development and disabling immunity, and is a comorbidity for several disorders including heart failure^4,5^.

Within cells, iron is utilised by ∼2% of proteins and ∼6.5% of enzymes (including ∼35% of oxidoreductases)^6^. Key conserved biochemical processes are iron-dependent, such as the electron transport chain (ETC), tricarboxylic acid (TCA) cycle, nucleotide synthesis, DNA repair, histone and DNA demethylation, and oxygen sensing^6^. Cells typically obtain iron by receptor-mediated internalisation of transferrin, a dedicated iron-chaperone protein in plasma and extracellular fluid^7^. However, the amount of iron bound to transferrin shows marked physiological variation. For example, in severe iron deficiency or in the presence of inflammation, plasma iron concentrations can drop by ∼90%^8,9^, drastically decreasing iron supply to cells.

Notably, human genetic studies and pre-clinical work link decreased transferrin-iron acquisition by proliferating lymphocytes to impaired adaptive immunity^5,10^. Patients homozygous for a mutation in *Tfrc* (the gene encoding the transferrin receptor, TFR1, also known as CD71) that inhibits cellular iron uptake, have a severe combined immunodeficiency characterised by reduced lymphocyte function and suppressed antibody titres^10^. Similarly, low serum iron availability suppresses B and T-cell responses in mice following vaccination and influenza infection^5^. How iron deficiency mechanistically impairs adaptive immune cells remains unclear, but understanding this issue could inform how iron regulates immunity and provide further rationale for correcting iron deficiency within human populations.

T-cells dramatically remodel their metabolism post-activation, including upregulation of glycolysis and oxidative phosphorylation (OXPHOS), in order to meet the increased energetic and biosynthetic requirements of proliferation and effector function^11^. Transition from naïve to activated T-cell is accompanied by greatly increased expression of the iron uptake protein, TFR1, and CD8+ T-cell iron content is calculated to triple within the first 24h post-activation^12^. In this study, we investigate how low iron supply influences activated CD8+ T-cells via ‘omics approaches and isotope tracing. We find mitochondrial defects, dysregulated metabolic processes and stalled cell division, but show that a single amino acid, aspartate, rescues multiple aspects of cell dysfunction including proliferation. These results offer new insight into the molecular effects of iron deficiency within the immune system.

## Results

### Transcriptomic and proteomic screens reveal potential nodes of dysfunction during CD8+ T-cell iron-deficiency

Iron is utilised by ∼400 proteins, of which 204 are described to be expressed in T-cells^6,12,13^. Iron interacting proteins operate in a wide diversity of pathways ranging from mitochondrial metabolism to DNA synthesis^6^, suggesting that iron-deficiency likely has important and wide-acting effects on T-cell biochemistry. However, the specific effects of iron scarcity are difficult to predict because of the interdependence of cellular processes. We therefore took an unbiased approach to initially define the impact of iron limitation on global CD8+ T-cell transcription and translation, employing RNA-sequencing (RNA-seq) and quantitative protein-mass spectrometry (protein-MS) of murine OT-I CD8+ T-cells activated *in vitro* across physiologically-titrated transferrin-iron concentrations (0.001-0.625 mg/mL holotransferrin, with total amount of transferrin kept constant at 1.2 mg/mL) for 48h (Fig. 1a).

**Fig. 1.**
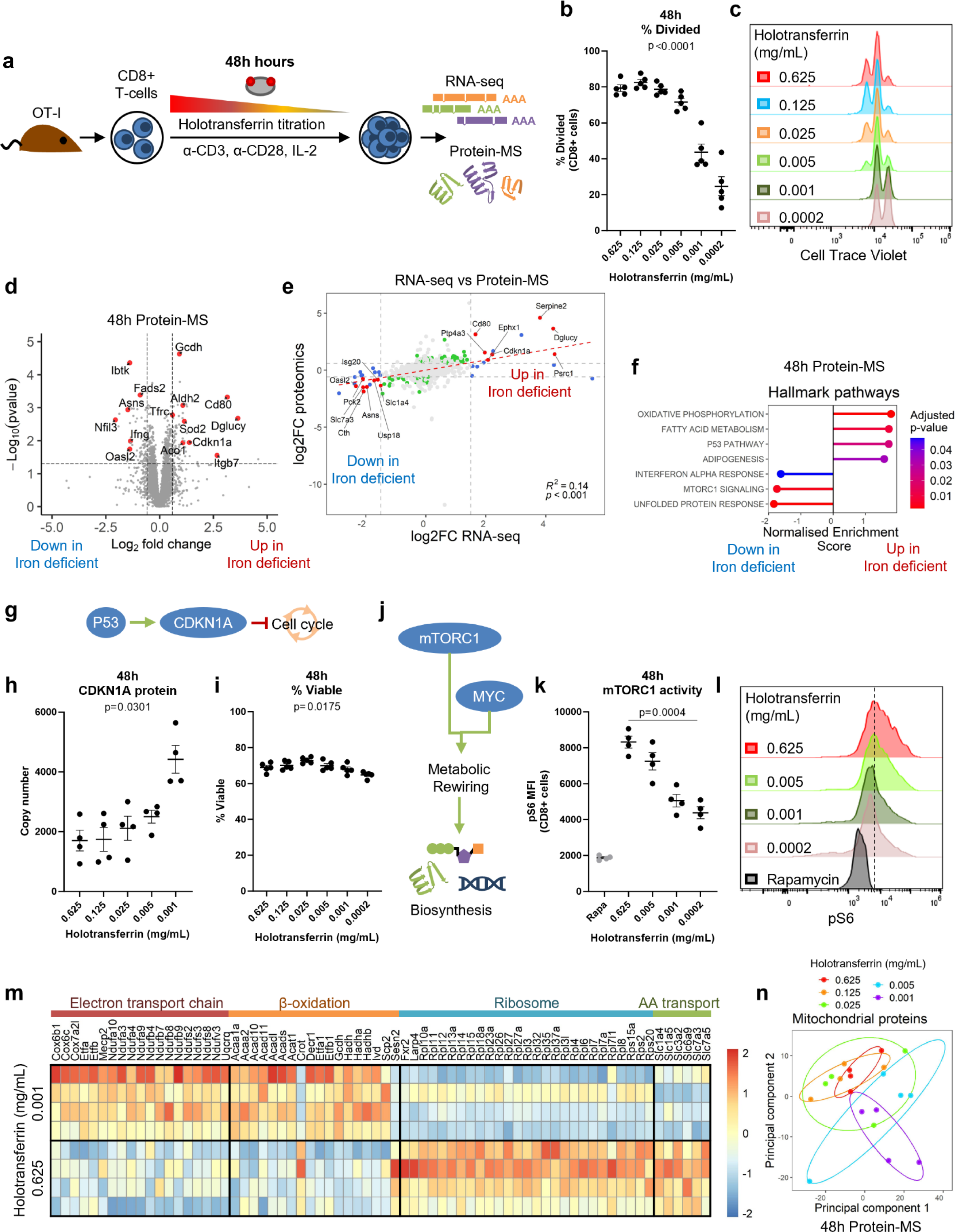
Iron-deficiency induces changes to metabolic processes at the RNA and protein level. **(a)** CD8+ OT-I T-cells isolated from mice were activated with 5 µg/mL plate bound α-CD3, 1 µg/mL α-CD28 and 50 U/mL IL-2 for 48h in a titration of holotransferrin conditions. Naïve T-cells for were collected on day 0. Where comparisons between high and low iron conditions are made, the holotransferrin concentrations used are 0.625 (high) and 0.001 (low) mg/mL. **(b-c)** Division assessed using CTV, n=5. **(d)** Volcano plot with the significance thresholds of log_2_|FC| > 0.585, p-value < 0.05, n=4. **(e)** Correlation plot of the log_2_|FC| between high and low iron conditions by RNA-seq and protein-MS, n=4. **(f)** Hallmark GSEA for the protein-MS, n=4. P53 induces CDKN1A expression which inhibits the G1-S phase transition. **(h)** CDKN1A protein expression and **(i)** percentage viable cells, n=4. **(j)** mTORC1 and MYC are metabolic regulators that enable biosynthesis downstream of TCR stimulation. **(k-l)** mTORC1 activity measured via its downstream target pS6, n=4. Controls were treated overnight with rapamycin (rapa; 1 µM). **(m)** Heatmap of proteins in selected metabolic pathways defined using GO terms where the p-value < 0.05, n=4. ETC = GO_RESPIRATORY_ELECTRON_TRANSPORT_CHAIN, β-oxidation = GO_FATTY_ACID_BETA_OXIDATION, ribosome = GO_CYTOSOLIC_RIBOSOME, amino acid (AA) transport = GO_AMINO_ACID_IMPORT. **(n)** PCA of the protein-MS given prior selection for proteins in the MitoCarta3.0 gene set, n=4. Data are mean ± SEM. Statistics are: **(a, h-i, k)** sampled matched one-way ANOVAs with the Geisser-Greenhouse correction; **(e)** Pearson correlation R^2^ value.

Despite observing suppression of cellular proliferation during iron deprivation, measured using cell trace violet (CTV) (Fig. 1b-c), only 193 genes were identified as significantly differentially regulated between iron replete and deficient conditions by RNA-seq (Supplementary Fig. 1a; Supplementary file 1). However, clear segregation was still observed between the two iron concentrations by principal component analysis (PCA; Supplementary Fig. 1b). Genes identified by RNA-seq as induced during iron-deficiency also displayed an enrichment of the activating histone marks, H3K4me3 and H3K27ac at transcription start sites (TSSs) in the low iron condition (Supplementary Fig. 1c) when analysed by ChIPmentation. Meanwhile, genes supressed by iron-deficiency at the RNA level showed depletion of these activating marks around their TSSs. This data indicates that RNA expression changes observed during iron deficiency are largely due to transcriptional differences rather than alterations in RNA stability. Expression of the iron uptake receptor, *Tfrc*, was significantly higher in the low iron condition (Supplementary Fig. 1d), consistent with cellular iron limitation^14^.

Protein-MS analysis identified 116 differentially expressed proteins (Fig. 1d; Supplementary file 2). As expected, TFR1 protein copy number increased as holotransferrin was depleted (Supplementary Fig. 1e). Despite the differential expression of 116 proteins, no differences were observed in overall protein mass or protein molecules per CD8+ T-cell activated across a titration of holotransferrin conditions (Supplementary Fig. 1f-g), indicating that despite decreased cell division, iron deficient CD8+ T-cells do not accumulate extra protein.

5091 mRNAs/proteins were mutually detected by the RNA-seq and protein-MS with a significant positive correlation in the log2|FC| values (Fig. 1e). Of these, only 15 were significantly differentially regulated at both the mRNA and protein levels (Supplementary Fig. 1h). These modest expression changes indicate changes in protein function rather than expression may be the predominant driver of the profound suppressive effect of iron deficiency on CD8+ T cell proliferation. Of note, 7 of the 15 mutually differentially expressed genes encoded proteins related to metabolism, including amino acid transporters (*Slc1a4*, *Slc7a3*) or enzymes (*Dglucy*, *Ephx1*, *Asns*, *Cth*, *Pck2*), indicating changes in cellular metabolism may underpin this functional alteration.

In addition, gene set enrichment analysis (GSEA), conducted on both RNA-seq and protein-MS datasets also indicated altered expression of the P53 pathway (Fig. 1f, Supplementary Fig, 1i). The P53 pathway coordinates responses to cellular stress such as DNA damage, hypoxia and nutrient deprivation, with downstream effects including apoptosis, DNA repair and cell cycle arrest^15^. Altered activity of this pathway may therefore also contribute to the titratable impairment to cellular proliferation in low iron conditions (Fig. 1b-c). Consistent with this, expression of CDKN1A, a P53 target and suppressor of the G1-S phase cell cycle transition was induced by iron deprivation at 24h, prior to the first cell division and continued to be upregulated as CD8+ T-cells entered their proliferative phase at 48h (Fig. 1g-h, Supplementary Fig. 1j-k). However, while cell division halted, cells remained viable at 48h (Fig. 1i). This data suggests that induction of the P53 signalling pathway may contribute to drive cell cycle suppression via CDKN1A, but iron deprivation does not cause cell death at this time point.

### Iron-deficiency does not induce a hypoxic response in CD8+ T-cells

P53 induction during iron chelation has been described in cell lines, macrophages and CD4+ T-cells^16–19^ and is understood to occur via HIF1α stabilisation due to impaired activity of their regulators, the iron and oxygen-dependent PHD proteins^19^. We therefore tested whether HIF1α was similarly stabilised in cells deprived of iron more physiologically, via decreased availability of transferrin-bound iron. While the iron chelator deferiprone (DFP) clearly induced HIF1α stabilisation in CD8+ T-cells, low iron conditions did not alter HIF1α levels relative to iron replete controls (Supplementary Fig. 1l-m). Moreover, the proteomic profile of iron deficient CD8+ T-cells bears little resemblance to the proteomes of 5 day activated cytotoxic CD8+ T-cells exposed to hypoxia for 24h as reported by Ross *et al*^20^ (Supplementary Fig. 1n). Both hypoxia and iron chelation have also been shown to induce glycolytic genes^16,20^, but this was not observed in our cells in low iron media.

### Low iron alters the CD8+ T-cell mitochondrial proteome

We observed decreased activity of the key upstream metabolic regulator, mTOR, measured as phosphorylation of its target S6 (pS6; Fig. 1j-l) and downregulation of the MYC and mTORC1 signalling pathways (Fig. 1f, Supplementary Fig. 1i). Decreased mTORC1 is consistent with lack of upregulation of glycolysis genes and may also relate to lack of total protein accumulation in non-proliferating iron deficient cells. However, unexpectedly, we observed upregulation of proteins involved in mitochondrial processes including OXPHOS and fatty acid metabolism (Fig. 1f). The enrichment of the fatty acid signature was driven largely by increases in β-oxidation proteins involved in fatty acid and branched chain amino acid breakdown rather than fatty acid synthesis (Fig. 1m). ETC proteins involved in the highly iron-dependent respiratory complexes I (CI), CIII and CIV were also upregulated (Fig. 1m). Consistent with increased abundance of these mitochondrial proteins, mitochondrial mass, measured with Mitotracker green (MTG), was also elevated in low iron conditions (Supplementary Fig. 1o). Of note, by selecting for mitochondrially localised proteins using the MitoCarta3.0 gene set^21^, we observed improved segregation within the first two principal components of CD8+ T-cells cultured in different iron concentrations (Fig. 1n) compared to when all proteins were analysed (Supplementary Fig. 1p), indicating mitochondrial proteins are disproportionately influenced by iron availability relative to all proteins detected.

Mitochondria are enriched for iron interacting proteins with 7% of mitochondrial proteins classified as iron-interacting compared to 2% cell wide^6^. Aconitase 2 (ACO2) and SDH of the TCA cycle and CI-CIV of the ETC require iron cofactors for function^22^. Mitochondria are also home to heme and iron-sulfur (Fe-S) cluster synthesis pathways^22^. Analysis of proteomics data from Howden *et al*^13^ revealed upregulation of heme and Fe-S cluster synthesis proteins in T-cells post-activation (Supplementary Fig. 2a-c). The heavy reliance of mitochondrial function on iron in combination with the upregulation of mitochondrial proteins suggests that mitochondrial function may be disrupted during iron scarcity.

### Iron-deficiency impairs CD8+ T cell mitochondrial function

To directly interrogate CD8+ T cell mitochondrial function under iron deprivation, we first assessed a metric of mitochondrial health, specifically levels of superoxide (O_2_^•–^) - a key exemplar of mROS species. As available iron declined, mROS levels increased in CD8+ T-cells (Fig. 2a-b). CI, CII and CIII of the ETC, which all require iron for electron transfer^22^ are key mROS producers^23^. Given the observed increase in ETC proteins (Fig. 1m), but lower availability of iron during iron restriction, we propose an imbalance in ETC proteins to iron cofactors may impair efficient electron transfer resulting in increased mROS generation. Consistent with this, iron deprivation decreased the mitochondrial membrane potential (Fig. 2c, Supplementary Fig. 3a) and was previously shown to suppress mitochondrial ATP generation^5^. Upregulation of the mitochondrial superoxide detoxifying protein, SOD2, was also observed under iron scarcity (Fig. 2d), indicating compensatory mechanisms to suppress mROS, which at the lowest iron concentrations still fail to control them.

**Fig. 2.**
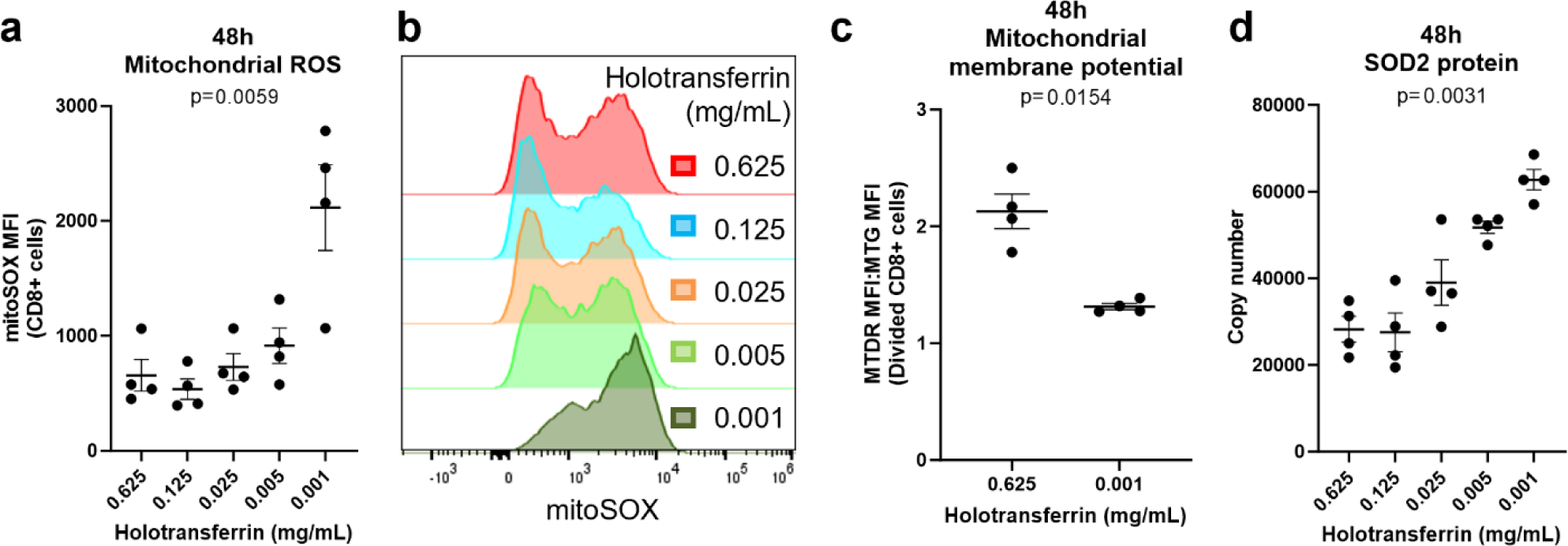
Iron deprivation induces mROS production and loss of mitochondrial membrane potential. CD8+ T-cells were activated as described in Fig. 1a. **(a-b)** mROS MFI measured using MitoSOX red, n=4. **(c)** Mitochondrial membrane potential calculated as the ratio of Mitotracker deep red (MTDR) to Mitotracker green (MTG), n=4. **(d)** SOD2 protein expression by protein-MS, n=4. Data are mean ± SEM. Statistics are: **(a, d)** matched one-way ANOVAs with the Geisser-Greenhouse correction; **(c)** matched two-tailed t-test.

Taken together with the alterations in the mitochondrial proteome, these data indicate that iron deprived CD8+ T-cells may have defective mitochondrial metabolism. To directly assess this, we next performed stable isotope-based tracing coupled to metabolite-mass spectrometry (metabolite-MS). CD8+ T-cells were activated for 24h and then incubated with ^13^C6-glucose or ^13^C5-glutamine for a further 24h under iron replete- or deficient conditions. Here, minimal changes in overall abundance of the glycolytic metabolites, pyruvate and lactate, or the amino acids derived from glycolytic intermediates (alanine, serine, and glycine) were observed, suggesting glycolytic activity was preserved under low iron conditions (Fig. 3a-b). This is consistent with minimal changes in HIF1α activity (Supplementary Fig. 1l-n) and in agreement with glycolysis being an iron independent pathway (none of the enzymes involved in glycolysis require iron).

**Fig. 3.**
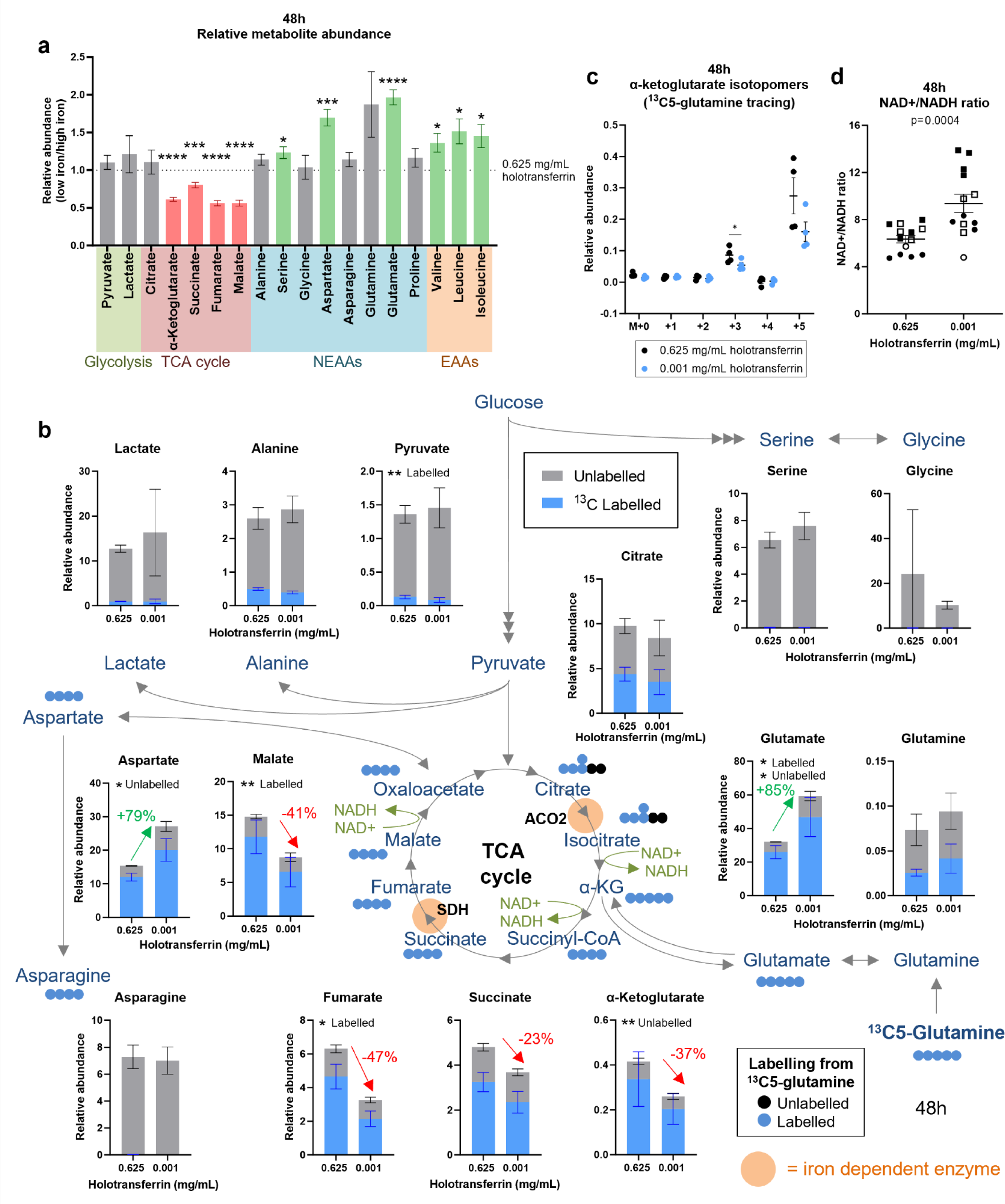
Iron scarcity impairs TCA cycle activity at the iron-dependent enzymes ACO2 and SDH. CD8+ T-cells were activated as described in Fig. 1a. For tracing experiments, T-cells were activated in standard media for 24h and then incubated in media containing ^13^C6-glucose or ^13^C5-glutamine for a further 24h. **(a)** Relative metabolite abundance from T-cells cultured in low iron (0.001 mg/mL holotransferrin) versus high iron (0.625 mg/mL holotransferrin) normalised to spiked in glutaric acid. Pooled relative total abundances from the ^13^C6-glucose and ^13^C5-glutamine experiments, n=8. NEAA = non-essential amino acids, EAA = essential amino acids. **(b)** ^13^C5-glutamine tracing, n=4. Relative abundance of labelled and unlabelled metabolites calculated as the fraction labelled multiplied by the raw abundance. **(c)** Relative abundance of α-KG mass isotopomers from ^13^C-glutamine tracing, n=4. **(d)** NAD+/NADH ratio. Data from independent experiments denoted by different symbols, n=13. Data are mean ± SEM. Statistics are: **(a)** matched t-tests between 0.625 and 0.001 mg/mL holotransferrin conditions for each metabolite; **(b, c)** matched two-way ANOVAs with the Geisser-Greenhouse correction and the Sidak correction for multiple comparisons; **(d)** matched two-tailed t-test. *p < 0.05, **p < 0.01, ***p < 0.001, ****p < 0.0001.

In these experiments we also noted that while ∼20% labelling of ^13^C6-glucose into the TCA cycle was observed, ^13^C5-glutamine was much more readily incorporated into the TCA cycle, labelling 60-80% of TCA cycle metabolites (Supplementary Fig. 3b). The larger incorporation of glutamine relative to glucose into the TCA cycle is consistent with reports indicating substantial reduction of glucose-derived pyruvate to lactate in activated T-cells, alongside increased glutamine anaplerosis into the TCA cycle^24^. The TCA cycle metabolite succinate lies upstream of an iron-dependent enzyme (SDH). Consistently, we observed that while succinate abundance was depressed in iron limiting conditions (−23%; Fig. 3a-b), it was relatively accumulated compared to the downstream metabolites, fumarate (−47%) and malate (−41%), suggesting suppressed flux from succinate to fumarate (Fig. 3a-b). Decreased ^13^C-labelling from ^13^C5-glutamine into fumarate and malate also supports this.

A second iron-dependent step of the TCA cycle occurs at the level of ACO2, which converts citrate to α-ketoglutarate (α-KG). In agreement with reduced TCA cycle activity, decreased abundance of α-KG was observed in low-iron cells (Fig. 3a-b). Decreased α-KG M+3 isotopomers from ^13^C5-glutamine and a trend towards reduced α-KG M+2 isotopomers from ^13^C6-glucose supports reduced ACO2 activity (Fig. 3c, Supplementary Fig. c-e). However, since α-KG is also produced from glutamate by activity of glutamate dehydrogenase (GDH), decreased α-KG may also reflect decreased activity of this enzyme. Indeed, decreased relative abundance of α-KG M+5 mass isotopomers from ^13^C5-glutamine, indicate a trend towards decreased GDH activity under low iron conditions (Fig. 3c, Supplementary Fig. 3d). Consistent with this, ^13^C-labelled glutamate accumulated in low iron conditions following ^13^C5-glutamine tracing (Fig. 3b). Decreased oxidative TCA cycle progression, due to limited glutamine anaplerosis, ACO2 and SDH activity agrees with diminished NAD+ reduction and consequent increases in the NAD+/NADH ratio (Fig. 3d, Supplementary Fig. 3f) also observed under iron scarcity. Since NADH is a positive allosteric regulator of GDH, this may also partly the indicated decrease in GDH activity^25^.

### Iron depletion alters H3K27 methylation, but proliferation is not rescued by α-KG supplementation

α-KG levels were decreased by ∼40% in iron-deficient CD8+ T cells (Fig. 3a-b, 4a). α-KG is a substrate for many dioxygenases, including the histone lysine demethylases (KDMs)^26^ (Fig. 4b) and alterations in T-cell metabolism have been shown to alter KDM activity and T-cell fate^27^. Crucially, most KDMs require an iron catalytic core to mediate hydroxylation of histone methyl groups using α-KG and oxygen as substrates^26^. The unstable methyl-hydroxy intermediate spontaneously dissociates leaving a demethylated product^28,29^. This dual dependency of KDMs for α-KG and iron mean that a decrease of α-KG availability under iron limitation may exert a double hit on KDM activity, with potential impacts on the appropriate chromatin restructuring necessary for CD8+ T-cell differentiation. The abundance of KDMs substantially increases upon T-cell activation, with KDM6B being the second most upregulated KDM in CD8+ T-cells^12,13^. KDM6B is responsible for the removal of the repressive histone mark, H3K27me3, a process critical for T-cell differentiation and effector function acquisition^30,31^. Consistent with decreased activity under iron limitation, CD8+ T-cells demonstrated a titratable failure to remove H3K27me3 relative to iron replete controls, with H3K27me3 levels remaining almost as high as IL-7 treated, “naïve-like” cells (Fig. 4c-d, Supplementary 4a-b). These findings therefore indicate that altered CD8+ T cell metabolic activity under low iron may have direct implications for their epigenetic status and differentiation capacity. Of note however, direct supplementation with cell-permeable dimethyl-α-KG failed to rescue H3K27me3 levels and cellular proliferation (Supplementary Fig. 4c-d) in iron deprived T-cells, consistent with H3K27me3 demethylation requiring both α-KG and iron.

**Fig. 4.**
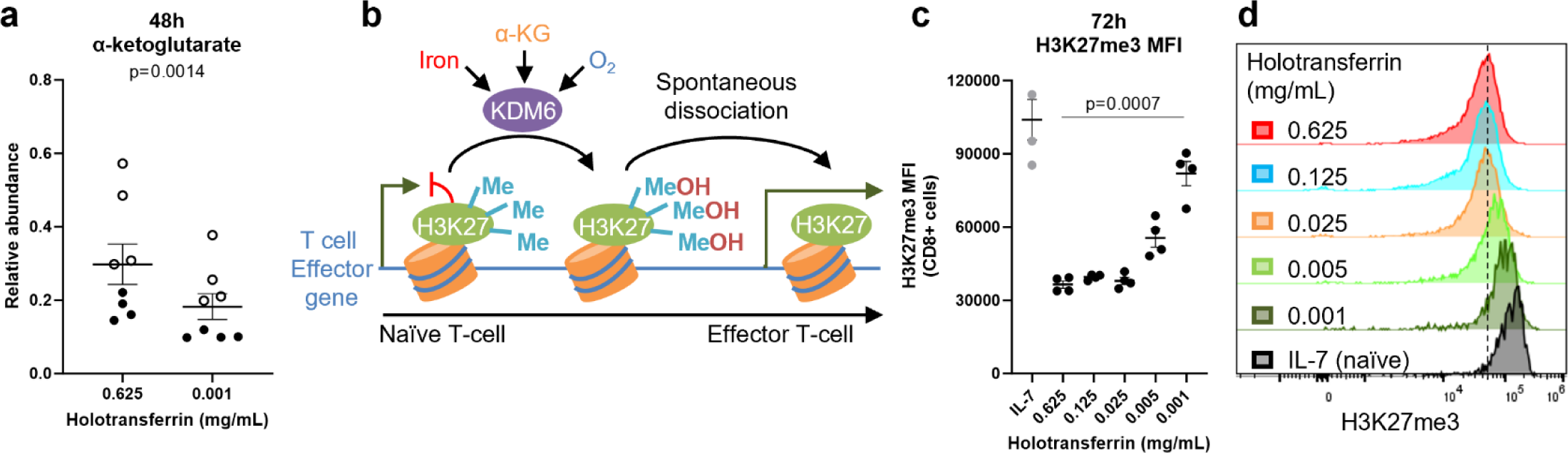
Iron-deficiency permits accumulation of H3K27me3 in CD8+ T-cells. CD8+ T-cells were activated as described in Fig. 1a. **(a)** Relative abundance of α-KG. Data from independent experiments denoted by different symbols, n=8. **(b)** KDM enzymes use iron cofactors and α-KG and oxygen substrates to mediate the hydroxylation of methyl groups which spontaneously dissociate to leave a demethylated histone. KDM6 enzymes remove the repressive histone mark, H3K27me3, from effector gene loci upon T-cell activation. **(c-d)** H3K27me3 MFI. “Naïve” controls cells were cultured in IL-7 (5 ng/mL), n=4. Data are mean ± SEM. Histograms are normalised to mode. Statistics are: **(a)** paired two tailed t-test; **(c)** matched one-way ANOVA with the Geisser-Greenhouse correction.

### Iron depletion suppresses nucleotide synthesis from aspartate

Aspartate is produced downstream of the TCA cycle metabolite oxaloacetate and critically supports cellular proliferation through de novo purine and pyrimidine synthesis^32–34^. In cancer cells, ETC inhibition or iron chelation impairs aspartate synthesis, resulting in suppressed proliferation of these immortalised cells^35–37^. However, despite TCA cycle inhibition, aspartate was unexpectedly higher in iron deficient cells, with the majority derived from glutamine (Fig. 3a-b, 5a, Supplementary Fig. 5a).

To understand whether nucleotide synthesis was altered downstream of decreased TCA activity in iron-depleted CD8+ T cells, we measured the abundance of nucleotides and their upstream precursors by LC-MS, observing that AICAR, a metabolite which lies two steps downstream of aspartate incorporation into purine synthesis^38^ was substantially decreased during iron limitation (Fig. 5b-c). PPAT, the initiating enzyme of purine synthesis which has been predicted to be iron dependent^6^ was also reduced in iron scarcity (Fig. 5d). Carbamoyl-aspartate and orotate which lie downstream of aspartate incorporation into pyrimidine synthesis were similarly depleted (Fig. 5e-f). Consistent with these observations, certain nucleotides were decreased during iron-deficiency (Supplementary Fig. 5b-c), albeit to a lesser extent, which may be explained by their decreased usage under suppressed CD8+ T-cell proliferation and associated DNA synthesis (Fig. 1b-c).

**Fig. 5.**
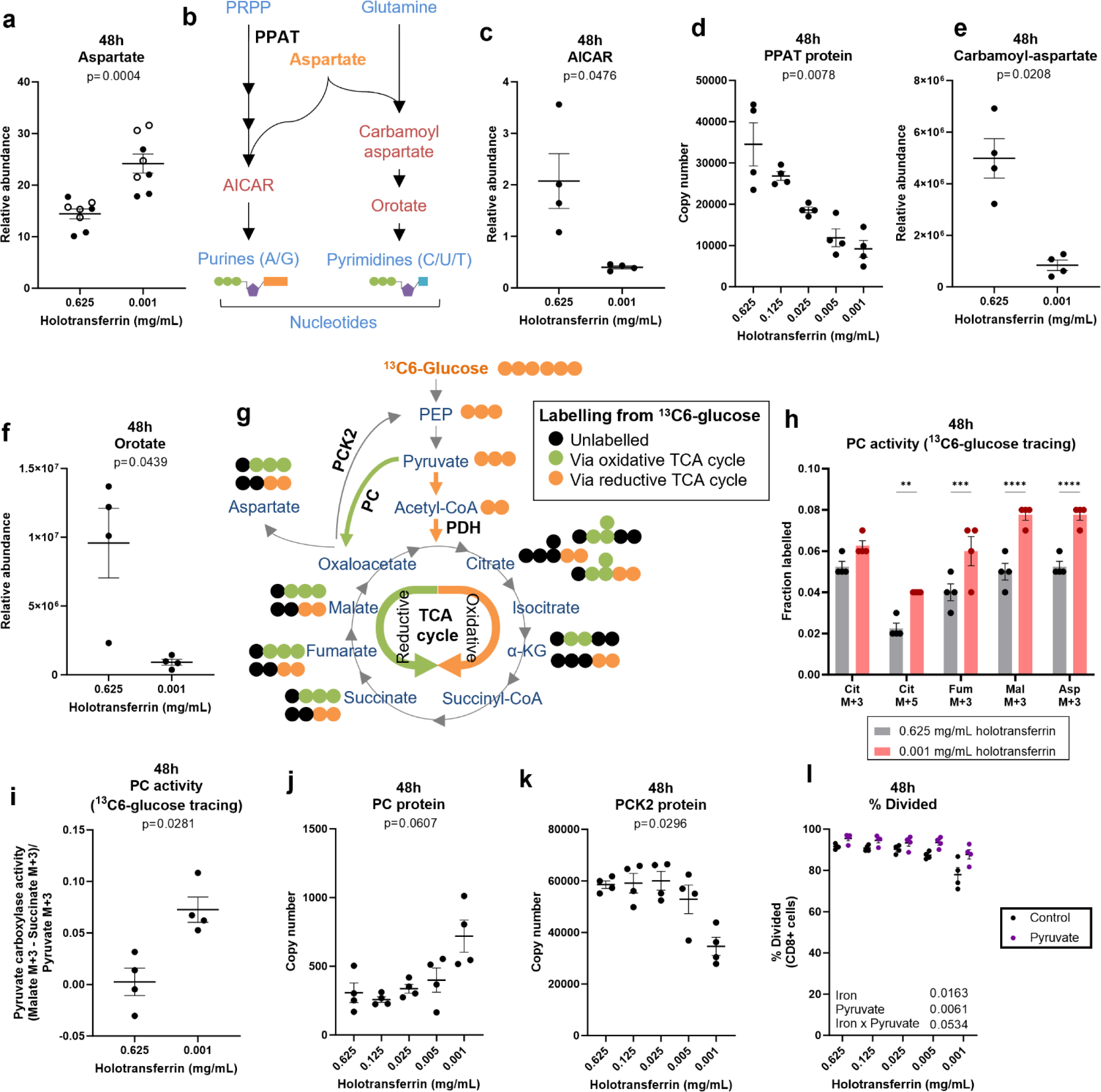
Iron scarcity suppresses nucleotide synthesis downstream of aspartate incorporation. CD8+ T-cells were activated as described in Fig. 1a. For the ^13^C6-glucose tracing experiments, T-cells were activated for 24h and then incubated in media containing ^13^C6-glucose for a further 24h. **(a)** Relative abundance of aspartate. Data from independent experiments denoted by different symbols, n=8. **(b)** Aspartate is incorporated into purine and pyrimidine nucleotides. Relative abundance of **(c)** AICAR, n=4. AICAR was normalised to spiked in ^15^N-dT. **(d)** PPAT protein via protein-MS, n=4. Relative abundance of **(e)** carbamoyl-aspartate and **(f)** orotate, n=4. Carbamoyl-aspartate and orotate were normalised to spiked in glutaric acid. **(g)** Schematic of ^13^C6-glucose tracing. Orange and green circles indicate ^13^C labelled atoms. Orange circles show labelling expected from oxidative TCA cycling via PDH. Green circles indicate labelling from reductive TCA cycling via PC. PC activity measured via the **(h)** fractional labelling into heavy labelled metabolites expected from reductive TCA cycling and via **(i)** the ratio of (Malate M3 – Succinate M3)/Pyruvate M3, n=4. **(j)** PC protein expression via protein-MS, n=4. **(k)** PCK2 protein expression via protein-MS, n=4. **(l)** Division measured using CTV with or without pyruvate (10 mM), n=4. Data are mean ± SEM. Statistics are: **(a, c, e-f, i)** paired two tailed t-tests; **(d, j-k)** one-way ANOVAs with the Geisser-Greenhouse correction; **(h)** matched two-way ANOVA with the Sidak correction for multiple comparisons; **(l)** matched two-way ANOVA with the Geisser-Greenhouse correction.

Together, these data indicate that nucleotide synthesis from aspartate is limited in iron deficient CD8+ T cells, despite overall cellular abundance of aspartate being increased rather than decreased (Fig. 3a-b, 5a). This aspartate accumulation could be partially explained by increased production by an alternative pathway, for instance pyruvate anaplerosis to oxaloacetate via pyruvate carboxylase (PC) and reductive TCA cycling (Fig. 5g)^39^. Carbon flux from ^13^C6-glucose through the oxidative (canonical forward) TCA cycle via pyruvate dehydrogenase (PDH) results in entry of two ^13^C atoms into TCA cycle metabolites (M+2 mass isotopomers)^39^. Meanwhile, entry of carbon from ^13^C6-glucose to the reductive (reverse) TCA cycle via PC produces M+3 mass isotopomers. During iron-deficiency, significant increases in ^13^C6-glucose labelling into M+3 TCA cycle metabolites including fumarate and malate were observed, indicative of increased PC contribution to these metabolites (Fig. 5h). PC activity, evaluated as (malate M+3 – succinate M+3)/pyruvate M+3, as utilised by Elia *et al*^40^, was also increased (Fig. 5i). PC protein also trended towards increased expression in low iron conditions (Fig. 5j) while PCK2, which mediates the reverse reaction, converting oxaloacetate to the glycolytic intermediate phosphoenolpyruvate (PEP) was suppressed (Fig. 5k) suggesting carbon flux into oxaloacetate is favoured in low iron conditions. In line with increased PC usage, pyruvate supplementation provided a proliferative advantage to iron depleted cells (Fig. 5l). A reversal of TCA cycle activity is also in agreement with the observed increase in the NAD+/NADH ratio (Fig. 3d, Supplementary Fig. 3f). This suggests that under iron limiting conditions, CD8+ T-cells may utilise PC and the reductive TCA cycle to replenish the depleted metabolites fumarate and malate, circumventing the use of the iron-dependent enzymes, ACO2 and SDH. Aspartate accumulation under iron depletion could also be explained by decreased usage, as shown for nucleotide synthesis above. Aspartate is also used for the synthesis of asparagine, however at 48h post-activation, CD8+ T-cells in both iron deficient and replete conditions had not synthesised asparagine, indicated by the absence of ^13^C labelling from ^13^C5-glutamine (Fig. 3b, Supplementary Fig. 5d). This agrees with previous reports that CD8+ T-cells lack asparagine producing capacity (asparagine synthetase expression) at 48h post-activation, but gain this later into their activation and differentiation^34^. Accordingly, asparagine supplementation provided no proliferative benefit (Supplementary Fig. 5e). Taken together, increased abundance of aspartate alongside decreased nucleotide synthesis indicates that aspartate usage by this pathway is suppressed under iron restriction.

### Increased nucleotide availability provides resistance to iron-deficiency in CD8+ T-cells

To interrogate whether nucleotide abundance limits CD8+ T cell proliferation under iron deficiency, we first tested whether genetically increasing nucleotide abundance could rescue proliferation. Cellular nucleotide balance is maintained via the activity of ribonucleotide reductase (RNR) and SAMHD1^41^ (Supplementary Fig. 6a). While RNR enables production of dNTPs, SAMHD1 degrades dNTPs^41^. Consistently, *Samhd1* deletion (KO) results in dNTP accumulation in lung fibroblasts^42^ and bone marrow derived DCs (BMDCs)^43^ and is assumed to operate similarly in CD8+ T-cells. We isolated CD8+ T-cells from *Samhd1*-KO or wildtype littermates and activated them as previously described. *Samhd1*-KO CD8+ T-cells were less sensitive to iron scarcity in terms of a block on proliferation relative to wildtype cells (Supplementary Fig. 6b). This effect was modest, likely explained by the fact that SAMHD1-KO prevents dNTP breakdown but cannot rescue impaired nucleotide production. SAMHD1-KO CD8+ T-cells showed appropriate upregulation of *Tfrc* expression in low iron concentrations (Supplementary Fig. 6c-d) but failed to rescue CD25 or perforin expression (Supplementary Fig. 6e-f). Notably, SAMHD1-KO cells showed comparable expression of *Cdkn1a* (Supplementary Fig. 6g), indicating increased nucleotide pools provide a proliferative advantage despite elevated *Cdkn1a* expression.

### Aspartate supplementation rescues iron deficient CD8+ T-cells

For aspartate to be utilised for nucleotide synthesis, it must be in the cytosol. Thus, an explanation for decreased aspartate usage could be inappropriate retention of aspartate in the mitochondria. SLC25A12/13 transports aspartate from the mitochondria to the cytosol in exchange for glutamate^44^ (Fig. 6a), but requires a proton gradient to mediate transport^44^, which is decreased in iron depleted CD8+ T-cells. (Fig. 2c). Therefore, it is possible that a reduction in ETC chain activity due to iron-deficiency may impair the mitochondrial proton gradient, in turn inhibiting SLC25A12/13 and sequestering aspartate in the mitochondria where it cannot be used for nucleotide synthesis.

**Fig. 6.**
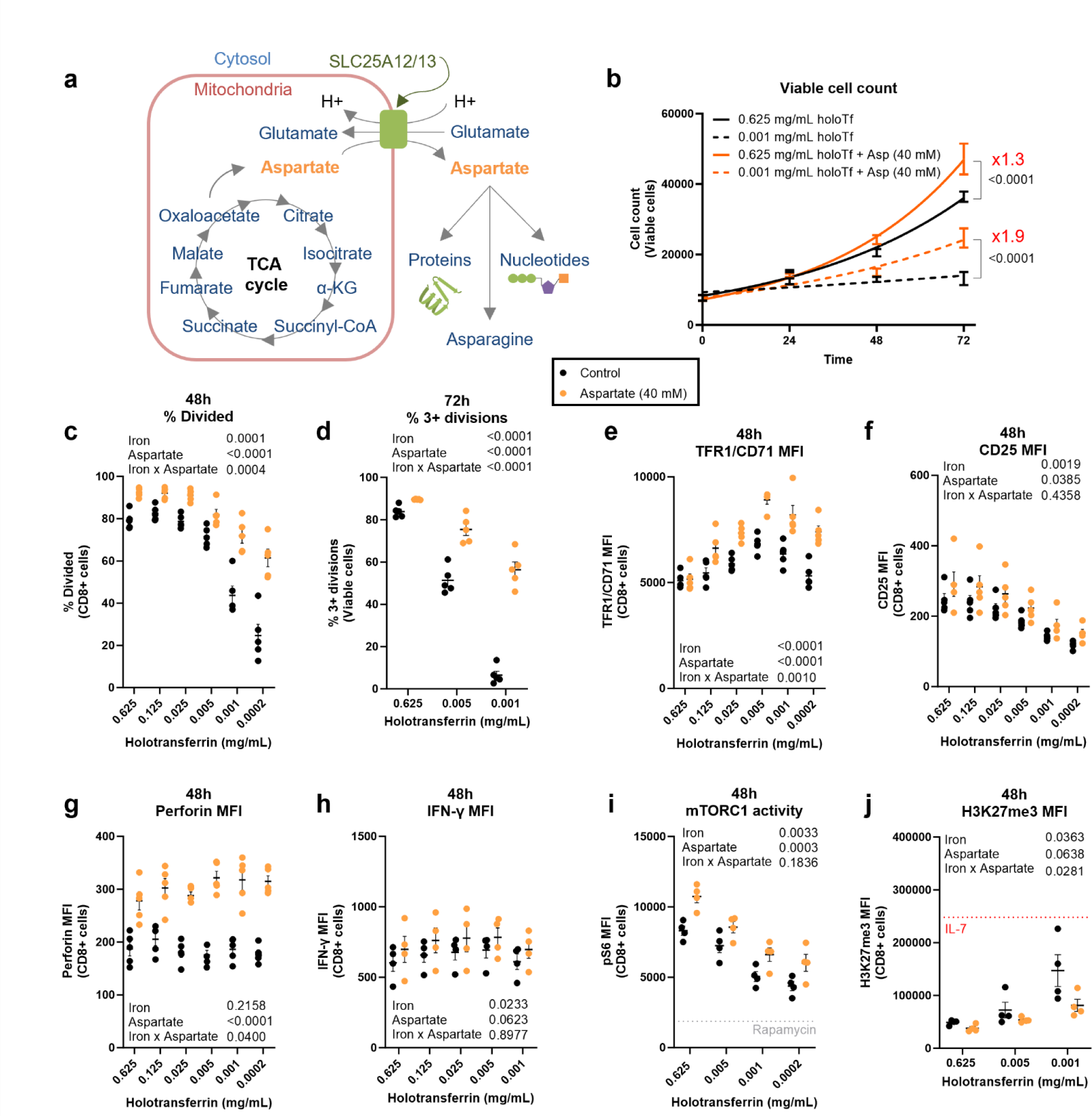
Aspartate increases the carrying capacity of iron deprived CD8+ T-cell cultures. CD8+ T-cells were activated as described in Fig. 1a with or without aspartate (40 mM). **(a)** Aspartate is synthesised in the mitochondria but must be transported into the cytosol by the proton-dependent transporter, SLC25A12/13 for downstream metabolism. **(b)** Viable cell counts, n=5. **(c)** Percentage divided cells at 48h and **(d)** percentage of cells undergoing 3+ divisions at 72h assessed using CTV, n=5. **(e)** TFR1/CD71**, (f)** CD25 and **(g)** perforin MFI, n=5. **(h)** IFN-γ MFI, n=4. **(i)** mTORC1 activity measured via pS6, n=4. **(j)** H3K27me3 MFI, n=4. “Naïve” controls cells were cultured in IL-7 (5 ng/mL). Data are mean ± SEM. Statistics are: **(b)** non-linear regressions using exponential growth equations with an extra sum-of-squares F test applied for either high or low holotransferrin concentrations between aspartate treated and untreated; **(c-j)** two-way ANOVAs with the Geisser-Greenhouse correction.

We directly assessed whether supplementation of aspartate into cell culture media, such that it can be taken up directly into the cytosol, can rescue CD8+ T cell proliferation. CD8+ T-cells were activated *in vitro* as described in Fig. 1a in the presence or absence of aspartate (40 mM). While CD8+ T-cells in low iron conditions with no added aspartate showed almost no population expansion over 72h of culture, the addition of aspartate increased the carrying capacity of the low iron culture by ∼10^4^ additional cells (Fig. 6b). When assessing division at 48h using CTV, aspartate increased the percentage of divided cells from 25% in the lowest iron condition (0.0002 mg/mL holotransferrin) to 61% (Fig. 6c). At 72h, the fraction of cells which could undergo three or more divisions in low iron conditions was profoundly increased with aspartate supplementation (increased from 7% to 56% in the lowest iron condition; Fig. 6d). Together, these data indicate that aspartate alone can substantially overcome the cell cycle impairment induced by iron scarcity and promotes a more profound recovery of cell division compared to exogenous pyruvate or S*amhd1*-KO.

While clonal expansion is critical for effective CD8+ T-cell effector responses, other aspects of CD8+ T-cell biology provide important indicators of function. *Tfrc* mRNA expression was increased in low iron concentrations, indicative of cellular iron scarcity, irrespective of whether aspartate was supplemented or not (Supplementary Fig. 7a). However, aspartate supplementation did permit CD8+ T-cells to maintain high TFR1/CD71 expression in the lowest iron concentrations when expression typically begins to drop (Fig. 7e). Aspartate may therefore provide benefit via promoting increased iron uptake capability during iron starvation. Aspartate supplementation also promoted CD8+ T cell expression of the activation marker, CD25, the cytolytic molecule, perforin, and the cytokine, IFN-γ, indicating its availability within the cytosol supports processes beyond proliferation (Fig. 6f-h).

To understand if aspartate, in addition to altering cell proliferation and function, induces a shift in gene expression, we conducted an RNA-seq of CD8+ T-cells in iron replete and iron deficient conditions, with or without aspartate (Supplementary file 3). Samples segregated largely based on iron concentrations rather than aspartate treatment (Supplementary Fig. 7b) suggesting aspartate supplementation does not drive broad alterations in transcriptional profile. Aspartate supplementation induced relatively few transcriptional changes with only 76 and 48 differentially expressed genes in high and low iron conditions respectively. Notably, the transcriptional changes induced by aspartate in low iron conditions correlate with the changes mediated by aspartate in high iron conditions (Supplementary Fig. 7c) indicating aspartate has similar, albeit small effects regardless of iron availability. Aspartate supplementation only marginally reduced *Cdkn1a* expression in low iron conditions (Supplementary Fig. 7d) indicating that the proliferative advantage conferred by aspartate is capable of overcoming P53 mediated cell cycle inhibition despite elevated *Cdkn1a*. GSEA revealed that the dominant aspartate effect is suppression of interferon (IFN) response pathways (Supplementary Fig. 7e) and was particularly driven by reduced IFN-stimulated gene-expression. Similar effects were observed for aspartate treatment at high iron concentrations (Supplementary Fig. 7f). Notably, type-I IFN can have anti-proliferative effects that may be related to upregulation of *Cdkn1a*^45^. Thus, the rescue effect of aspartate on CD8+ T-cell division may be partly facilitated if anti-proliferative type-I IFN signalling is counteracted.

Aspartate did not alter mROS generation (Supplementary Fig. 7g), suggesting that aspartate is unable to attenuate the mitochondrial dysfunction likely occurring at CI and CIII in iron deprived CD8+ T-cells. However, aspartate did increase mTORC1 activity measured via the expression of the downstream target, pS6 (Fig. 6i) and slightly suppressed the iron-deficiency-mediated increase in the NAD+/NADH ratio (Supplementary Fig. 7h). Aspartate treated CD8+ T-cells also displayed increased glycolytic and total ATP production (Supplementary Fig. 7i-k) and counteracted the accumulation of the repressive histone mark H3K27me3 by iron deficient CD8+ T-cells (Fig. 6j). Therefore, even in cells cultured in very low iron conditions, aspartate availability enables TCR-triggered CD8+ T-cells to acquire an activated phenotype, reconfigure important aspects of their metabolism, and proliferate.

## Discussion

Given the breadth and conserved nature of pathways involving iron interacting proteins it follows that iron deprivation would likely have broad implications on cell biochemistry, health, and function with potential knock-on effects at the tissue or systemic levels. Our work explains why iron depletion potently suppresses T-cell responses in models of immunisation, infection, and autoimmunity^5,17,46–48^. We also propose that the iron deficiency-associated perturbations described here may in part be responsible for cellular dysfunction in other cell types and tissues during iron deficiency.

Cells which undergo rapid proliferation are more likely to be impacted by iron-deficiency than quiescent populations. This is because in iron scarce environments, dividing cells will dilute out iron stores between daughter cells without exogenous iron pools to replenish the difference, and dividing cells have high biosynthetic demands. Meanwhile, quiescent cells can furnish iron interacting proteins via internal recycling of iron cofactors. This principle is demonstrated by T-cells homozygous for the TFR1^Y20H^ mutation which reduces iron uptake efficiency^10^. Quiescent naïve TFR1^Y20H^ T-cells are not noticeably impacted by the mutation but become extremely sensitive upon activation^10^. Moreover, some tissues, such as muscle may be able to resist iron deficiency to some degree by upregulation of glycolysis^49^. However, the capacity to switch to a glycolytic program may depend on having reduced demands for nucleotides and other macromolecular synthesis processes necessary for proliferation.

We demonstrate that iron-deficiency in CD8+ T-cells promotes an altered metabolic state. Iron-deficiency caused relatively modest perturbation of the cellular transcriptome and proteome, at odds with the significant inhibition of proliferation. Iron-deficiency acts most immediately to derail metabolism, with the most drastic effects on mitochondrial function. We observed decreased TCA cycle activity at the iron-dependent enzymes ACO2 and SDH, resulting in depletion of downstream metabolites. Despite suppressed TCA cycling, aspartate levels were augmented, but metabolites immediately downstream of aspartate metabolism, used for cytosolic nucleotide synthesis, were suppressed. We propose that aspartate accumulates within mitochondria due to iron-deficiency associated suppressed activity of the proton-dependent mitochondrial aspartate carrier, SLC25A12/13. Remarkably, aspartate supplementation of culture media, assumed to access the cytosol directly, profoundly rescued iron-deficiency impaired proliferation and enhanced other functional aspects of activated T-cells including H3K27me3 removal and increased mTORC1 activity. Our results place disrupted aspartate handling front and centre in the metabolic derangements of cellular iron-deficiency.

Increased H3K27me3 in iron deficient CD8+ T-cells is suggestive of KDM6 dysfunction. The inability of CD8+ T-cells to remodel their chromatin environment likely has implications for cellular differentiation. Moreover, suppressed levels of α-KG and/or iron may be capable of inhibiting iron and α-KG-dependent enzymes more generally including other KDM enzymes, the ten-eleven translocation (TET) DNA demethylases, as has been seen for KDM3B, which can act as an iron sensor influencing H3K9 methylation and mTORC1 activation^50^, and the PHD proteins which regulate HIF1α^19,51^. However, iron scarcity did not stabilise HIF1α or induce a proteomic response resembling that induced under hypoxic conditions. Different iron-dependent enzymes may have differential responses to iron inaccessibility, likely due to different iron binding affinities and expression levels. The PHD proteins have very high affinities for iron^52^, potentially explaining our result that iron chelators which potently suppress iron availability induce a HIF1α response while iron deficiency caused by low extracellular transferrin-iron supply cannot. Thus, iron deprivation does not blanket inhibit all iron-dependent pathways and iron chelation does not fully recapitulate physiological iron-deficiency.

In 1925, Otto Warburg proposed that “every living cell contains iron and that life without iron is impossible”^53^, referring to the fact that cell culture media requires iron in order to support cellular growth^53^. Activated iron deficient T-cells do not progress through the proliferative cycle, but also do not apoptose, as has been observed by others^50^. Cellular iron deficiency mediated by depriving physiological sources of extracellular iron imparts a distinct and dysregulated metabolic signature. Our findings demonstrate that a lack of iron alters mitochondrial function, with knock-on effects influencing utilisation of cytosolic and nuclear metabolic pathways, ultimately affecting cell identity. Iron deficient activated CD8+ T-cells are ‘stunned’ but not moribund, and can be substantially restored by aspartate, partially overcoming Warburg’s limitation. Aspartate is generally poorly taken up by cells, but our findings suggest that engineering cells, such as tumour targeted chimeric antigen receptor T-cells (CAR-T cells), to overexpress the plasma membrane aspartate transporter (SLC1A1/2/3) could provide resistance to iron depletion, for example in niches such as the tumour microenvironment.

## Acknowledgements

Thank you to the staff of the Department of Biomedical Services, University of Oxford for animal husbandry and welfare, the members of the Drakesmith lab, Michael Murphy (MRC Mitochondrial Biology Unit, University of Cambridge), Sumana Sharma (MRC Translational Immune Discovery Unit, University of Oxford), Katja Simon (The Kennedy Institute of Rheumatology, University of Oxford) and Ana-Victoria Lechuga-Vieco (The Kennedy Institute of Rheumatology, University of Oxford) for helpful discussions, Natasha White and Giulia Pironaci (both MRC Translational Immune Discovery Unit, University of Oxford) for assistance with OT-I colony management, Hannah Murray and Dana Costigan (both MRC Translational Immune Discovery Unit, University of Oxford) for practical help with pilot experiments, Andrew Howden and the FingerPrints Proteomic Facility for protein-MS analysis (University of Dundee).

M.R.T, J.N.F., A.E.P., A.E.A and H.D. were supported by a U.K Medical research council, MRC Human Immunology Unit grant awarded to H.D. (MCU_12010/10). M.R.T. is funded by the Clarendon Fund and the Corpus Christi College A. E. Haigh graduate scholarship. T.A.M. and A.S. were supported by a Medical Research Council (MRC, UK) Molecular Haematology Unit grant awarded to T.A.M. (MC_UU_00016/6 and MC_UU_00029/6). L.V.S was supported by a Wellcome Trust Principal Research Fellowship, awarded to Doreen Cantrell (205023/Z/16/Z). N.G. and S.D. are supported by a Blood Cancer UK Project grant (21007) and an MRC new investigator research grant (MR/V011588/1). J.R., B.M. and D.A.T are supported by a Cancer Research UK Programme grant (C42109/A24757).

## Author contributions

Conceptualisation, M.R.T., H.D., S.D., J.N.F., A.E.A., L.V.S., B.K.D., A.L.S., D.A.T, A.E.P., S.J.D., T.A.M., J.R.; Methodology, M.R.T., N.G., L.V.S, A.L.S.; Formal analysis, M.R.T.; Investigation, M.R.T, N.G., J.N.F., L.V.S., A.L.S., C.L.M., B.K.D., J.R., B.M.; Resources, J.R.; Writing, M.R.T, H.D., A.E.A.; Supervision, H.D., A.E.A., S.K.D.; Funding acquisition, H.D., S.D.

## Data availability

The RNA-seq and ChIPmentation datasets have been submitted to the Gene Expression Omnibus (GEO) with the dataset identifiers (GSE251962, GSE251963, GSE251964). The protein-MS dataset has been deposited to the ProteomeXchange Consortium via the PRIDE partner repository with the dataset identifier (PXD047814)

## Methods

### Mice

Animal work was completed under the authority of UK home office project and personal licenses granted under the Animals (Scientific Procedures) Act (ASPA) 1986. Mice were housed in individually ventilated cages. OT-I CD45.1 mice were acquired from Vincenzo Cerundolo, University of Oxford and Audrey Gerard, University of Oxford. *Samhd1*-KO mice and their genotyping were described in Rehwinkel et al^43^. C57BL6/J mice were purchased from Envigo. Mice were euthanised using a rising concentration of CO_2_ followed by cervical dislocation.

### Cell culture media

To manipulate iron availability, iron free medium was prepared with RPMI 1640 media (Gibco, 21875034), 1% penicillin/streptomycin (Sigma Aldrich, P0781-100ML), 1% glutamine (Sigma Aldrich, G7513-100ML) and 10% iron free serum substitute (Pan Biotech, P04-95080). Iron free media was supplemented with set concentrations of human holotransferrin (R&D systems, 2914-HT-001G/Sigma Aldrich, T0665) ranging from 0.0002-0.625 mg/mL. Total transferrin was kept constant at 1.2 mg/mL by adjusting human apotransferrin (unbound transferrin) concentrations (R&D systems, 3188-AT-001G/Sigma Aldrich, T1147) accordingly. The highest concentration of 0.625 mg/mL holotransferrin (∼15.6 µmol/L of iron) is reflective of the levels of iron physiologically found in human sera (14-32 µmol/L of iron^54^). Meanwhile, the low holotransferrin concentrations of 0.001 mg/mL are much lower than that found in plasma, but are hypothesised to be representative of highly iron depleted environments such as those in the tumour microenvironment^37^.

Aspartate (Scientific Laboratory supplies, CHE2306) was dissolved in iron free media by slowly titrating in 1M NaOH (Sigma Aldrich, 30620-1KG-M) until aspartate was completely solubilised and the pH ∼7.5. Asparagine and pyruvate were dissolved directly into iron free media at the described concentrations.

### CD8+ T-cell activation

Plates for CD8+ T-cell activation were coated with 5μg/mL α-CD3 (Biolegend, 100239) in phosphate buffered saline (PBS) for 2-3h at 37°C. Lymph nodes and spleens (inguinal, axillary, brachial, cervical, and mesenteric) from mice were sterilely collected in iron free media and homogenised through 40 μm filters using EasySep buffer (Stem Cell Technologies, 20144) or an in-house alternative (PBS + 2% FBS + 1mM EDTA (Invitrogen, AM9260G)) to produce single cell suspensions. CD8+ T-cells were isolated from total homogenate using the EasySep Mouse CD8+ T-cell (Stem Cell Technologies, 19853) isolation kit and EasyPlate EasySep magnets (Stem Cell Technologies, 18102) or EasyEights EasySep magnets (Stem Cell Technologies, 18103), according to the manufacturer’s protocols. Cells were optionally stained with cell trace violet (CTV, Invitrogen, C34557) for 8 minutes at 37°C prior to culture. Cells were plated at 0.5-1×10^6^ cells/mL in iron free media supplemented with defined holotransferrin concentrations (see section cell culture media). Cells were provided with 50 μM β-mercaptoethanol (BME, Gibco, 31350-010), 1 μg/mL α-CD28 (Biolegend, 102115) and 50 U/mL IL-2 (Biolegend, 575402). CD8+ T-cells were cultured at 37°C, 5% CO_2_ for 24-72h.

### RNA extraction

CD8+ T-cells were collected from cell culture plates and washed twice with PBS; cell pellets were immediately snap frozen on dry ice. T-cell pellets were resuspended in 350 µL of RLT+ buffer from the Qiagen RNeasy plus mini kit (Qiagen, 74136) and RNA was extracted according to the manufacturer’s instructions. For RNA extracted for RNA-sequencing, samples were treated on-column with the Qiagen RNase-free DNase I set (Qiagen, 79254) according to the method described in appendix D of the Qiagen RNeasy mini kit manual (Qiagen, 74104). RNA concentrations were determined using a Nanodrop one instrument (Thermofisher Scientific).

### cDNA synthesis and qPCR

cDNA was synthesised using the High-Capacity RNA-to-cDNA kit (Applied Biosystems, 4388950) and qPCR experiments were completed on a QuantStudio 7 flex real-time PCR system (Applied Biosystems, 4485701) using the Taqman gene expression master mix (Applied Biosystems, 4369016) and appropriate Taqman Gene Expression Assay (*B2m,* Mm00437762_m1; *Cdkn1a,* Mm04205640_g1; *Tfrc,* Mm00441941_m1). *B2m* was used as the endogenous control gene.

### RNA-sequencing

For RNA-sequencing, RNA quality was assessed using an Agilent high sensitivity RNA ScreenTape (Agilent, 5067-5579) and corresponding sample buffer (Agilent, 5067-5580) on a 4200 TapeStation system (Agilent, G2991BA). Library preparation and bulk mRNA-sequencing to a depth of 30 million paired end reads using the Illumina NovaSeq 6000 platform was conducted by Novogene.

Read alignment was conducted using RNA-star to the *Mus Musculus* genome (mm39) and features were annotated using FeatureCounts from the subread package. Differential gene expression analysis was conducted using EdgeR (R package) with the thresholds of log_2_|fold change| > 1.5 and an FDR <0.05 applied. Gene set enrichment analysis (GSEA) was completed using FGSEA (R package) and Hallmark pathways.

### ChIP-mentation

CD8+ T-cells were activated for 48h after which they were collected, washed twice in PBS, and fixed in 1% paraformaldehyde (PFA, Thermo Scientific, 28906) for 10 minutes at room temperature with rotation. Cells were washed twice at 16000 g for 30 seconds and dry pellets were snap frozen on dry ice.

Antibody binding buffer was prepared: 0.5% BSA and 1/200 protease inhibitor cocktail (PIC) in PBS. 10 µL of protein A dynabeads (ThermoFisher, 1001D) per sample were washed twice with antibody binding buffer on a magnet. Beads were blocked with 1 µL of antibody (α-H3K27ac, Diagenode, C15410196; α-H3K4me3, Diagenode, C15410003) in antibody binding buffer for 3-4h at 4°C with rotation. Just prior to adding the chromatin, the beads were washed with antibody binding buffer.

Cell pellets were resuspended in 120 µL of lysis buffer (50 mM Tris pH8, 10 mM EDTA, 0.5% SDS) with 1/200 PIC and transferred into Covaris tubes (Covaris, 520045). Samples were sonicated for 180 seconds using the settings aiming for 200 bp fragments. Samples were diluted with lysis buffer with 1/200 PIC to a 900 µL total volume and 100 µL of 10% Triton X-100 was added to neutralise. Samples were incubated for 10 minutes with rotation. 10 µL of prewashed protein A dynabeads were added and incubated for 30 minutes at 4°C. Samples were placed on a magnet and 250 µL of sample supernatant was added to antibody bound beads. Chromatin/antibody/bead mixtures were incubated overnight at 4°C with rotation. Samples were washed three times with 150 µL RIPA wash buffer (50 mM HEPES pH 7.6, 500 mM LiCl, 1 mM EDTA, 1% NP-40, 0.7% NaDeoxycholate), once with 150 µL TE buffer and once with 150 µL 10 mM Tris pH 8, each time for 5 minutes. Beads were resuspended in 29 µL pre-warmed tagmentation buffer with 1 µL Tn5 transpose (Illumina, 20034197). Samples were incubated for 5 minutes with vigorous mixing at 1100 rpm at 37°C in a thermomixer. 150 µL of RIPA buffer was added immediately to degrade the Tn5. Beads were washed once with 150 µL of RIPA buffer and then resuspended with 22.5 µL of ddH_2_O.

25 µL of 2X NEBNext Ultra II Q5 MasterMix (New England Biosciences, M0544S), 1.25 µL 5 µM universal adaptor primer and 1.25 uL 5 µM index adaptor primer was added to each sample. Samples were transferred to a thermocycler and the following program was run: 72°C for 5 minutes then 95°C for 5 minutes followed by 12 cycles of 98°C for 10 minutes, 63°C for 30 seconds and 72°C for 3 minutes. Following the 12 cycles, samples were incubated at 72°C for 5 minutes and then held at 12°C. Supernatants were removed and 50 µL of Ampure XP beads (Beckman Coulter, A63880) were added and pipetted 10 times and then incubated at room temperature for 2 minutes. Beads were washed once with 150 µL 80% ethanol, incubating for 2 minutes at room temperature. The ethanol was removed, and the beads were allowed to air dry for 3-5 minutes. 11 µL of ddH_2_O was added to the beads to elute the DNA. The supernatant was collected.

DNA quality was assessed using an Agilent D100 high sensitivity screentape (Agilent, 5067-5584) and the High sensitivity D1000 reagents (Agilent, 5067-5585) on a 2200 TapeStation system (Agilent, G2964AA) and using a Qubit fluorometer. DNA-sequencing to a depth of 10 million paired end reads using the Illumina NovaSeq 6000 platform was conducted by Novogene.

Data was analysed using the SeqNado analysis pipeline (https://github.com/alsmith151/SeqNado). Peak enrichment for H3K27ac and H3K4me3 was conducted for the promoters of genes previously identified as being differentially regulated at the RNA level by the RNA-seq.

### Protein-mass spectrometry

CD8+ T-cells were activated for 48h, harvested, and washed twice in PBS. Cells were fixed in 2% PFA (Thermo Scientific, 28906) for 30 minutes, washed and resuspended in PBS. Cells that were alive at the time of fixation were sorted by flow cytometry using forward and side scatter into PBS. Cells were pelleted and snap frozen on dry ice. Pelleted cells were lysed using the 2-step trypsin lysis protocol described by Kelly et al^55^. Cell pellets were resuspended in 200 µL TEAB digest buffer (0.1 M TEAB, 1 mM MgCl2, 1:80 benzonase, pH 8) and incubated for 20 minutes at 37°C in a thermomixer. The amount of trypsin for a 1:20 trypsin to protein (w/w) ratio was calculated and 50% of the required trypsin was added. Samples were incubated overnight at 37°C in a thermomixer. The remaining trypsin was added, and samples were incubated for 60 minutes at 37°C in a thermomixer. The samples were acidified to a final concentration of 1% trifluoroacetic acid (TFA) and subjected to a C18 stage-tip desalting with the following buffers: condition (100% acetonitrile), wash (0.1% TFA), elute (66.6% acetonitrile, 0.1% TFA), ion exchange (2:1 ratio of 100% acetonitrile to 0.1% TFA). Samples were dried using a speed-vac at 65°C.

Protein-MS analysis was performed as described previously^56^. Peptides were analysed on a Q-Exactive-HF-X (Thermo Scientific) mass spectrometer coupled with a Dionex Ultimate 3000 RS (Thermo Scientific). LC buffers were the following: buffer A (0.1% formic acid in Milli-Q water (v/v)) and buffer B (80% acetonitrile and 0.1% formic acid in Milli-Q water (v/v)). 2 μg of each sample were loaded at 15 μL/min onto a trap column (100 μm × 2 cm, PepMap nanoViper C18 column, 5 μm, 100 Å, Thermo Scientific) equilibrated in 0.1% TFA. The trap column was washed for 3 minutes at the same flow rate with 0.1% TFA then switched in-line with a Thermo Scientific, resolving C18 column (75 μm × 50 cm, PepMap RSLC C18 column, 2 μm, 100 Å). The peptides were eluted from the column at a constant flow rate of 300 nl/minute with a linear gradient from 3% buffer B to 6% buffer B in 5 minutes, then from 6% buffer B to 35% buffer B in 115 minutes, and finally to 80% buffer B within 7 minutes. The column was then washed with 80% buffer B for 4 minutes and re-equilibrated in 3% buffer B for 15 minutes. Two blanks were run between each sample to reduce carry-over. The column was always kept at a constant temperature of 50°C.

The data were acquired using an easy spray source operated in positive mode with spray voltage at 1.9 kV, the capillary temperature at 250°C and the funnel RF at 60°C. The MS was operated in data-independent acquisition (DIA) mode as reported earlier with some modifications^57^. A scan cycle comprised a full MS scan (m/z range from 350–1650, with a maximum ion injection time of 20 ms, a resolution of 120,000 and automatic gain control (AGC) value of 5 × 10^6^). MS survey scan was followed by MS/MS DIA scan events using the following parameters: default charge state of 3, resolution 30.000, maximum ion injection time 55 ms, AGC 3 × 106, stepped normalized collision energy 25.5, 27 and 30, fixed first mass 200 m/z. Data for both MS and MS/MS scans were acquired in profile mode. Mass accuracy was checked before the start of sample analysis.

Quantification of reporter ions was completed using Spectronaut (Biognosys; Spectronaut 14.10.201222.47784) in library-free (directDIA) mode. Minimum peptide length was set to 7, and maximum peptide length was set to 52, with a maximum of 2 missed cleavages. Trypsin was specified as the digestive enzyme used. The FDR at the precursor ion level and protein level was set at 1% (protein and precursor Q value cutoff). The max number of variable modifications was set to 5, with protein N-terminal acetylation and glutamine and asparagine deamidation and methionine oxidation set as variable modifications. Carbamidomethylation of cysteine residues was selected as a fixed modification. Data filtering and protein copy number quantification were performed in the Perseus software package, version 1.6.6.0. Copy numbers were calculated using the proteomic ruler^58^. This method sets the summed peptide intensities of the histones to the number of histones in a typical diploid cell. The ratio between the histone peptide intensity and summed peptide intensities of all other identified proteins is then used to estimate the protein copy number per cell for all the identified proteins. Data was subsequently analysed using custom R scripts. A log_2_|fold change| > 0.585 (equivalent to |fold change| > 1.5) was used (as used in Howden et al^13^). We also applied the typical p-value threshold of <0.05 when comparing the high (0.625 mg/mL holotransferrin) and low (0.001 mg/mL holotransferrin) iron concentrations via a t-test as well as an additional threshold of p-values < 0.05 when a one-way ANOVA was conducted across all conditions. For RNA-seq, GSEA was conducted using FGSEA (R package) and the Hallmark pathways. For analysing mitochondrial proteins, proteins were filtered by inclusion in the MitoCarta3.0 gene set^21^. Data Availability: The mass spectrometry proteomics data have been deposited to the ProteomeXchange Consortium via the PRIDE^59^ partner repository with the dataset identifier PXD047814.

### ^13^C6-glucose and ^13^C5-glutamine tracing

For heavy isotope tracing experiments, CD8+ T-cells were isolated and activated as described above. Iron free media with varying holotransferrin concentrations was prepared using phenol red free RPMI 1640 (Gibco, 11835030) as phenol red can interfere with metabolite-ms. At 24h prior to cell collection (24h post activation), cells were collected, washed, and replated in media containing the heavy isotope of interest (13C6-glucose (Cambridge Isotope Laboratories, Inc., CLM-1396-1) and 13C5-glutamine (CK Isotopes, CNLM-1275)) on plates coated with α-CD3. Iron free tracing media was prepared using SILAC RPMI 1640 flex media (Gibco, A24945201) which lacks glucose, phenol red, glutamine, arginine, and lysine. Argine (MP Biomedicals, 194626) and lysine hydrochloride (MP Biomedicals, 194697) were supplemented at standard RPMI 1640 concentrations of 1.2 mM and 0.2 mM respectively. For 13C6-glucose tracing, glutamine (Sigma Aldrich, G7513-100ML) was added at 2 mM and 13C6-glucose at 11.1 mM. For 13C5-glutamine tracing, 13C5-glutamine was added at 2 mM and glucose (Sigma Aldrich, 158968-100G) at 11.1 mM. Holotransferrin and apotransferrin as well as activation reagents (BME, α-CD28 and IL-2) were added to the iron free tracing media as described in section 2.3 at standard concentrations. Metabolites were measured by Gas chromatography-mass spectrometry as described below.

### Gas chromatography-mass spectrometry (GC/MS) for metabolites

48h activated CD8+ T-cells were collected, counted and 2-4×10^6^ cells were washed twice in ice cold 0.9% saline made with NaCl (Sigma Aldrich, 31434-500G-M) and ultrapure HPLC grade water (Alfa Aesar, 22934). Cells were pelleted and snap frozen on dry ice. Cells were extracted in 1:1:1 pre-chilled methanol, HPLC-grade water (containing 1.75 μg/mL D6-glutaric acid) and chloroform. The extracts were shaken at 1400 rpm for 15 minutes at 4°C and centrifuged at 12,000 g for 15 minutes at 4°C. The upper aqueous phase was collected and evaporated under vacuum. Metabolite derivatization was performed using an Agilent autosampler. Dried polar metabolites were dissolved in 15 μL of 2% methoxyamine hydrochloride in pyridine (Thermo Fisher Scientific, 25104) at 55°C, followed by an equal volume of N-tert-Butyldimethylsilyl-N-methyltrifluoroacetamide with 1% tertbutyldimethylchlorosilane after 60 minutes, and incubation for a further 90 minutes at 55°C. GC-MS analysis was performed using an Agilent 6890GC equipped with a 30m DB-35 MS capillary column. The GC was connected to an Agilent 5975C MS operating under electron impact ionization at 70 eV. The MS source was held at 230°C and the quadrupole at 150°C. The detector was operated in scan mode and 1 μL of derivatised sample was injected in splitless mode. Helium was used as a carrier gas at a flow rate of 1 mL/min. The GC oven temperature was held at 80°C for 6 minutes and increased to 325°C at a rate of 10°C/minutes for 4 minutes. The run time for each sample was 59 minutes. For determination of the mass isotopomer distributions (MIDs), spectra were corrected for natural isotope abundance. Data processing was performed using MATLAB.

### Liquid chromatography-mass spectrometry (LC/MS) for AICAR detection

CD8+ T-cells were collected after 48h of culture and 3×10^6^ cells were washed twice with PBS. Dry pellets were snap frozen on dry ice. Cell pellets were resuspended in 0.01% formic acid (Sigma Aldrich, 1003445799) containing 0.8 µM 15N_2_-dT (Cambridge Isotope Labs, NLM-3901-PK) and were incubated for 37°C for 30 minutes. Samples were centrifuged at maximum speed for 10 minutes and the lysates were collected and added to 3 kDa amicon ultra 0.5 centrifugal filter units (Millipore, UFC500324) and centrifuged for 20 minutes at 16000 g. The filtrates were transferred to a mass spectrometry vial for analysis of compounds in nucleotide synthesis. Samples were analysed on a TSQ Altis Triple Quadrupole Mass Spectrometer in selected reaction monitoring mode (SRM) interfaced to an UltiMate 3000 uHPLC. The uHPLC was fitted with a nanoEase M/Z Symmetry C18 Trap Column, 100Å, 5 µm, 180 µm × 20 mm (Waters) at RT and a Luna Omega 3µm PS C18 column (150 x 0.3 mm) connected to an EASY-Spray™ source with an EASY-Spray Cap flow emitter (15µM). 10 µl of sample was injected per run via a 10 µl sample loop. Buffers used were from Romil and of Ultra LC standard. Buffer A: H_2_O (0.1 % acetic acid), buffer B MeCN (0.1 % acetic acid). The gradient was 0-2.8 min – 1 % B, 22 min – 30 % B, 23.5 min – 99 % B. This was followed by 2 wash pulses (1-99 % B) and equilibration to 1% B (45 min total run time). The trap column was held at a constant 1 % B and switching from the trap to the main column occurred at 1 min and back at 40 min. Mass spectrometry conditions were as follows: source voltage of 2400V in positive ionisation mode; ion transfer tube temperature 275 °C, CID gas pressure 1.5 mTorr, scan widths for Q1 and Q3 at 0.7 *m/z*, a chromatographic filter was used with a peak width of 6 sec, dwell time 20 ms. Collision energy voltage and RF voltage were optimized using the Thermo Tune software using for the compounds assayed using authentic compounds dissolved in Buffer A. Peaks retention times were confirmed in samples by co-injection with the relevant standard.

Data was analysed using the FreeStyle 1.6 software and the Genesis peak detection algorithm with its default settings. Data was analysed by ratioing the target peak area with the ^15^N_2_-dT peak area.

### Liquid chromatography-mass spectrometry (LC/MS) for nucleotides and nucleotide precursors

48h CD8+ T-cells were collected at 48h post-activation and 3×10^6^ cells were washed twice with ice cold 0.9% saline made with NaCl (Sigma Aldrich, 31434-500G-M) and ultrapure HPLC grade water (Alfa Aesar, 22934). Cells were pelleted and snap frozen on dry ice. Metabolites were extracted with 100 µL of ice-cold extraction buffer (40% methanol (Biosolve, BIO-13687802), 40% acetonitrile (Biosolve, BIO-01204102), 20% water (Biosolve, BIO-23214102-1), 15 µM glutaric acid internal standard and 0.5% formic acid (Biosolve, BIO-069141A8)) for 5 minutes on ice. 8.8 µL of 15% ammonium bicarbonate (Supelco, 5.33005) solution was added to neutralise and samples were left on dry ice for a further 15 minutes. Samples were thawed on ice and then centrifuged at 20 G at 4°C for 5 minutes. Supernatants were collected for LC-MS analysis.

LCMS analysis was performed on an Agilent LCMS QToF 6546 using a Waters Premier BEH Z-HILIC VanGuard Fit column (1.7 µm, 2.1 mm x 150 mm) with the following mobile phases: Mobile phase A: 20 mM Ammonium hydrogen carbonate (LiChropur, Supelco) in LC-MS grade water (Ultra CHROMASOLV, Honeywell Riedel-de Haën) with 0.1% ammonium hydroxide (Alfa Aesar, Thermo Scientific) and 5 µM InfinityLab deactivator additive (Agilent Technologies).

Mobile phase B: 90% UHPLC grade acetonitrile (BioSolv, Greyhound Chemicals), 10% LC-MS grade water (Ultra CHROMASOLV, Honeywell Riedel-de Haën) with 5 uM InfinityLab deactivator additive (Agilent Technologies).

5 µl of sample was injected and the chromatographic separation was achieved with a gradient run with a constant flow rate of 0.2 mL/min and the following program: T = 0 min, 10% A, 90% B; T = 2 min, 10% A, 90% B; T = 18 min, 35% A, 65% B; T = 22 min, 65% A, 35% B; T = 22.1 min, 85% A, 15 % B; T = 25 min, 85% A, 15% B; T = 25.1 min, 10% A, 90% B; T = 30 min, 10% A, 90% B.

Full scan data was acquired between m/z 50 – 1050 at 1 Hz whilst using online mass correction. Analyte ionisation was achieved via ESI (negative polarity) with the following parameters: VCap: 2000 V, Nozzle Voltage: 500 V, gas temperature: 225 °C, drying gas: 8 l/min, nebulizer: 30 psi, sheath gas temp: 300 °C, sheath gas flow 12 l/min. Data extraction was performed using Agilent Profinder 10.0. Data was normalised to the internal standard (D6-Glutaric acid) peak area using an in-house R script.

### NAD+/NADH ratio measurements

NAD+/NADH ratio measurements were made using the NAD/NADH-Glo assay (Promega, G9071) according to the manufacturer’s instructions with modifications for measuring NAD+ and NADH individually (rather than pooled). Cultured CD8+ T-cells were collected, washed in PBS, and counted. 1×10^5^ cells in 50 µL of PBS were lysed in 50 µL of 0.2 M NaOH (Sigma Aldrich, 30620-1KG-M) with 1% dodecyltrimethylammonium bromide (DTAB, Alfa Aesar, A10761) per condition in technical duplicates. Each sample was split (50 µL) into a second tube for matched NAD+ and NADH measurements.

For measurement of NAD+, NADH was first degraded by adding 25 µL of 0.4 M HCl (Fisher Scientific, H/1111/PB17) to the first tube, heated to 60°C for 15 minutes followed by incubation at room temperature for 10 minutes. 25 µL 0.5 M TRIZMA base (Sigma Aldrich, T1503-1KG) was added to quench the HCl.

To measure NADH, NAD+ was degraded by heating the second tube to 60°C for 15 minutes followed by incubation at room temperature for 10 minutes. 50 µL of 0.2 M HCl (Fisher Scientific, H/1111/PB17) and 0.25 M TRIZMA base (Sigma Aldrich, T1503-1KG) solution was added to quench the NaOH.

The NAD/NADH-Glo detection reagent was prepared as instructed. 50 µL of each sample (either purified for NAD+ or NADH) and 50 µL of NAD/NADH-Glo detection reagent was added to a white 96 well plate and mixed gently. The plate was incubated at room temperature and luminescence was measured using a Promega GloMax multi-detection luminometer at 30, 45 and 60 minutes.

### Extracellular flux analysis assay

Extracellular flux analysis (Seahorse) was conducted using the Seahorse XF Real-Time ATP rate assay kit (Agilent, 103592-100). The day prior to the assay, the Agilent Seahorse XF96 analyser was turned on and a sensor cartridge (Agilent, 103792-100) was hydrated in sterile dH2O at 37°C in a non-CO_2_ incubator overnight. The water was replaced by Seahorse XF calibrant (Agilent, 103059-000) on the day of the assay and the cartridge was incubated for 1 hour prior to loading the drug ports. The day of the assay, ATP assay media was prepared using 1 mM pyruvate (Agilent, 103578-100), 2 mM glutamine (Agilent, 103579-100), 10 mM glucose (Agilent, 103577-100) in Seahorse XF RPMI pH 7.4 (Agilent, 103576-100) and warmed to 37°C. Cells were collected, washed and 1×10^5^ cells in 50 µL of ATP assay media were plated per well on a Poly-D-lysine (Gibco, A3890401) coated XF96 microplate (Agilent, 101085-004). Plates were spun down, supernatant removed, and cells incubated at 37°C for 30 minutes in a non-CO_2_ incubator followed by addition of 130 µL of ATP assay media and further incubation for 25-30 minutes. Oligomycin and rotenone/antimycin A were used at a final concentration of 1.5 µM and 0.5 µM respectively. 20 µL of oligomycin (15 µM) and 22 µL of rotenone/antimycin A (5 µM) were loaded into the sensor cartridge in ports A and B respectively. The ATP rate assay was run on a Seahorse XF96 analyser with 3 measurements taken for each step (baseline, oligomycin, rotenone/antimycin A) using the following injection settings: 3 minutes mix, 0 minutes wait, 3 minutes measure. Data was analysed using the Agilent Seahorse Wave desktop software (version 2.6) and the Seahorse XF real-time ATP rate assay report generator.

### Flow cytometry

Cells were transferred from cell culture plates to 96 well round bottom plates and washed with PBS.

In intracellular cytokine staining experiments, cells were first stimulated with cell activation cocktail (1:500 or 1:2500) (Biolegend, 423301), BFA (5 mg/mL) (Biolegend, 420601) and monensin (2 mM) (Biolegend, 420701) in iron free media at 37°C/5% CO_2_ for 5h prior to staining.

Cells were stained with 20 µL of surface antibody cocktail with Zombie NIR fixable viability kit (1:400-1000, Biolegend, 423105) in PBS for 20 minutes on ice. Cells were fixed with 2% paraformaldehyde (Pierce, 28906) diluted in PBS or commercial fixation buffer (Biolegend, 420801) for 20 minutes on ice. For nuclear staining of markers such as H3K27me3, cells were fixed in Foxp3 transcription factor fixation buffer (eBioscience, 00-5523-00) for 1h on ice.

For intracellular staining, cells were permeabilised in perm/wash buffer (Biolegend, 421002) for 20 minutes. Intracellular targets were stained with 20 µL of intracellular stain prepared in perm/wash buffer for 20 minutes to overnight. Cells were resuspended in PBS and data was acquired using an Attune NxT flow cytometer (Thermofisher Scientific). Data was analysed using FlowJo (BD biosciences).

Detection of mROS was conducted using MitoSOX dye (Invitrogen, M36008). Cells were resuspended in 200 µL of MitoSOX dye diluted to 5 µM in phenol red iron free media and incubated at 37°C/5% CO_2_ for 15 minutes before surface staining. Acquisition was conducted on live cells.

Cells were stained with Mitotracker Green (MTG; M7514, Thermofisher Scientific) and Mitotracker Deep Red (MTDR; M22426, Thermofisher Scientific) at 100nM in iron free media for 30 minutes at 37°C/5% CO_2_ prior to staining. Cells were acquired live. The ratio of MTDR to MTG was calculated as a metric of mitochondrial membrane potential relative to mitochondrial mass.

For flow staining for the metabolic regulators, pS6 and HIF1α, cells were washed once with a solution of 50% RPMI 1640, 50% HBSS (Gibco, 14025-092) and 0.5% BSA (PAN biotech, PO6-139310) before fixation in 1% PFA (Pierce, 28906) for 20 minutes. Cells were washed with PBS and optionally surface stained for fixation stable epitopes. Cells were permeabilised using 90% ice cold methanol (Sigma Aldrich, 34860-1L-R) for 20 minutes at −20°C prior to permeabilization and acquisition.

### Cell counting by flow cytometry

Cells were transferred into 96 well round bottom TC plates and washed once with 2% FBS in PBS (FACS buffer). Cells were resuspended in FACS buffer containing 0.5 µg/mL 7-AAD (Biolegend, 420403) and incubated on ice for 10 minutes and then acquired directly on the Attune NxT flow cytometer (Thermofisher Scientific). 50 µL of cells were acquired per well to enable accurate calculation of cell concentration and wells were resuspended using a multichannel between each column on the 96 well plate to account for cell settling effects.

### Data analysis

Unless explicitly specified, data analysis was completed using Excel (Microsoft), Prism (GraphPad), FlowJo (BD biosciences) or custom scripts written in the R coding language. Statistics are specified in Fig. legends.

---

**Supplementary Fig. 1.**
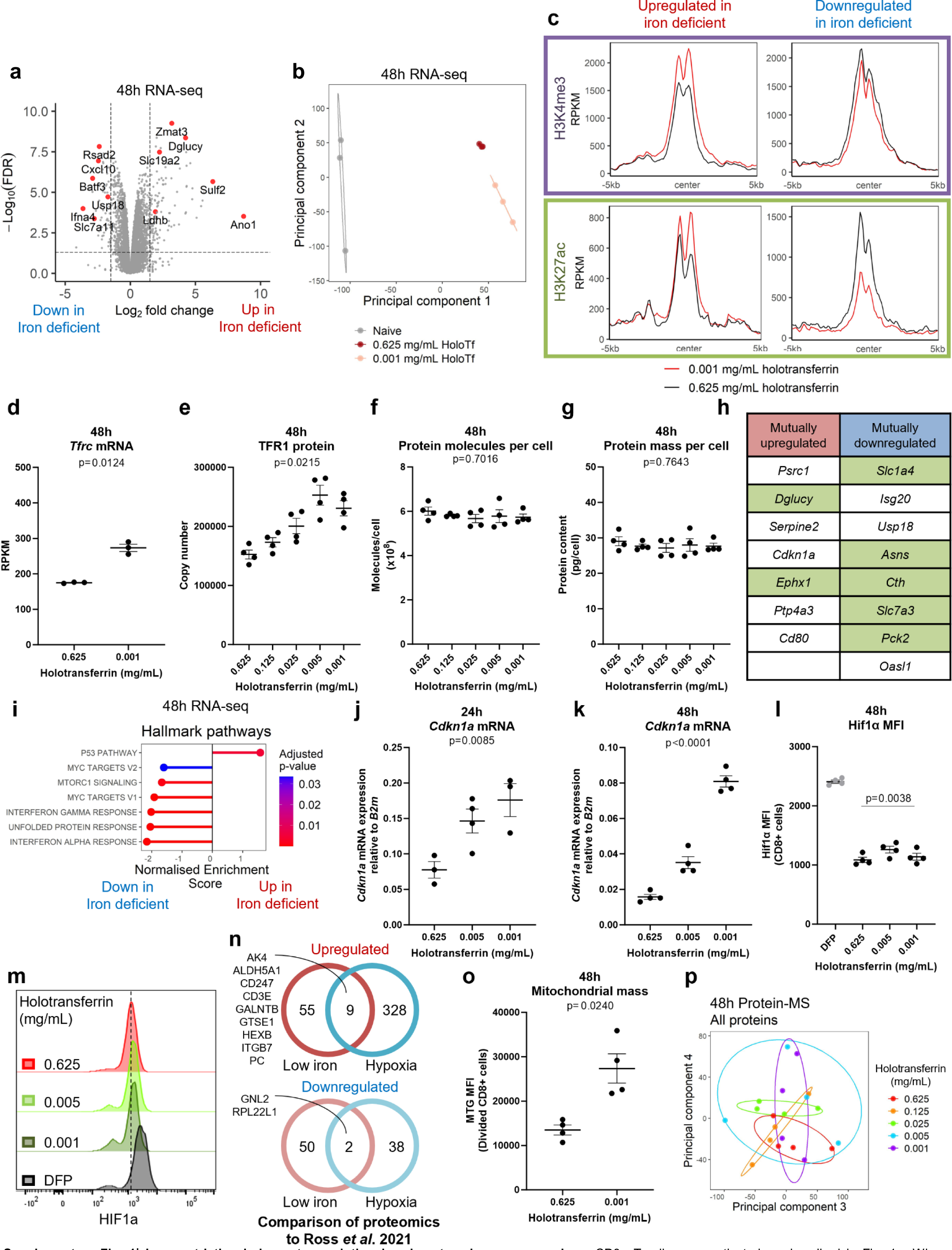
Iron restriction induces transcriptional and proteomic reprogramming. CD8+ T-cells were activated as described in Fig. 1a. Where comparisons between high and low iron conditions are made, the holotransferrin concentrations used are 0.625 (high) and 0.001 (low) mg/mL. **(a)** Volcano plot of RNA-seq with significance thresholds of FDR < 0.05 and log_2_|FC| > 1.5, n=4. **(b)** PCA of RNA-seq samples, n=4. **(c)** H3K4me3 and H3K27ac enrichment at TSSs of genes identified to be up or downregulated by iron-deficiency, n=1. Center = TSS. RPKM = reads per kilobase of transcript per million mapped reads. *Tfrc*/TFR1 **(d)** mRNA and **(e)** protein expression, n=4. **(f)** protein molecules and **(g)** protein mass per cell measured via protein-MS, n=4. Genes and proteins mutually up or downregulated in low iron conditions, n=4. Metabolic genes are in green. **(i)** Hallmark GSEA for the RNA-seq, n=4. *Cdkn1a* mRNA expression assessed by qPCR at **(j)** 24h and **(k)** 48h, n=4. **(l-m)** HIF1α MFI, n=4. Controls were treated for 4h with deferiprone (DFP; 75 µM). **(n)** Venn diagrams of proteins mutually up or downregulated in response to iron deficient conditions and hypoxia (1% O_2_ relative to normoxia at 18% O_2_) in CD8+ T-cells. Hypoxia data are derived from Ross *et al*^20^. **(o)** Mitochondrial membrane mass measured using Mitotracker green (MTG) MFI, n=4. **(p)** Principal components analysis of all proteins, n=4. Statistics are: **(d, o)** paired t-test; **(e-g, j-l)** sample matched one-way ANOVAs with the Geisser-Greenhouse correction.

**Supplementary Fig. 2.**
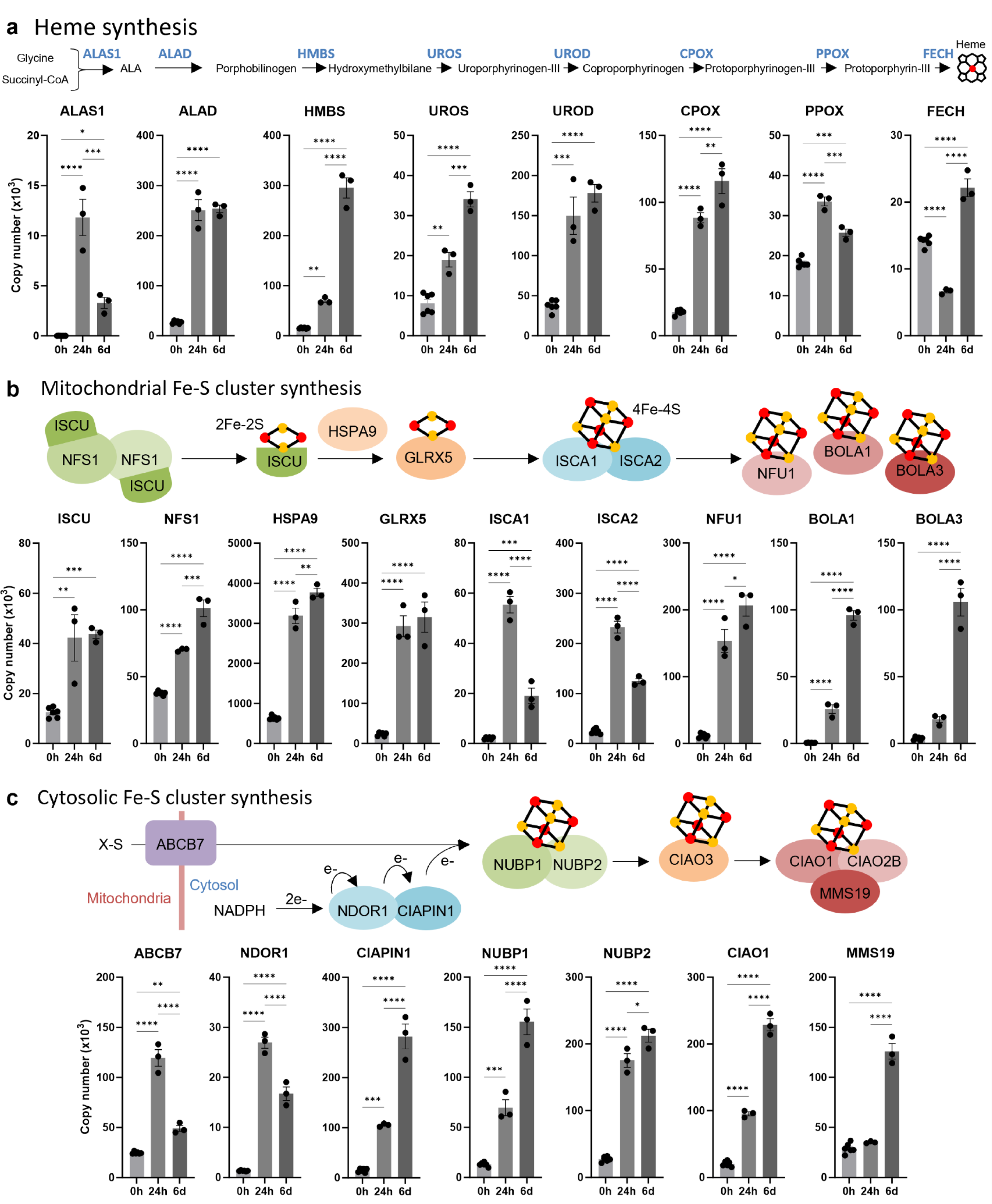
Heme and Fe-S cluster synthesis proteins are induced following CD8+ T cell activation. Data derived from Howden *et al*^13^. Protein copy numbers for proteins involved in **(a)** heme synthesis, **(b)** mitochondrial Fe-S cluster synthesis and **(c)** cytosolic Fe-S cluster synthesis. n=6 at 0h, n=3 at 24h and 6d. Data are mean ± SEM. Statistics are: **(a-c)** ordinary one-way ANOVAs with multiple comparisons using Tukey’s correction. *p < 0.05; **p < 0.01; ***p<0.001; ****p < 0.0001.

**Supplementary Fig. 3.**
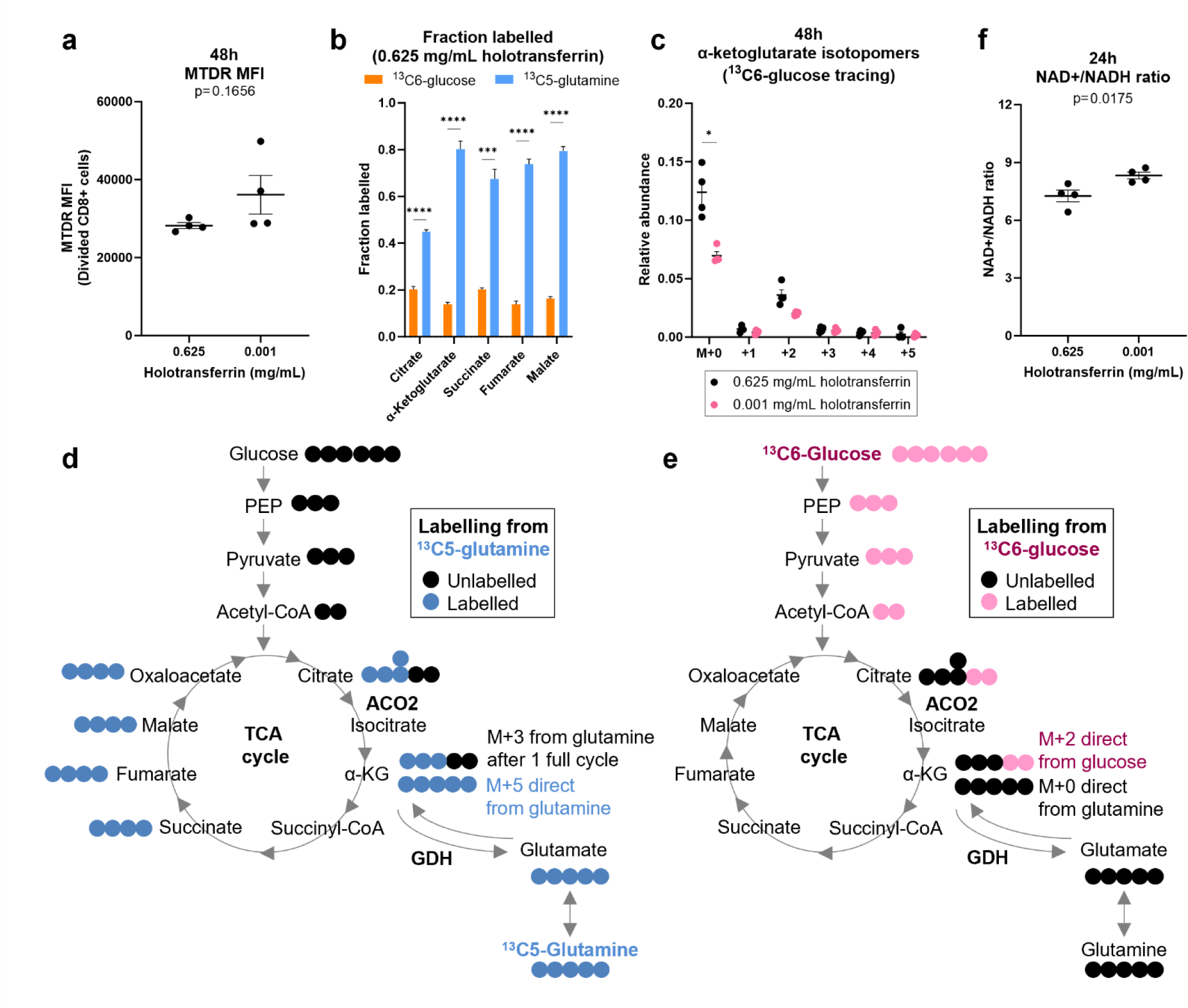
Iron-deficiency alters CD8+ T-cell mitochondrial metabolism. CD8+ T-cells were activated as described in Fig. 1a. **(a)** Mitotracker deep red (MTDR) MFI, n=4. **(b)** Metabolic fraction labelled by either ^13^C6-glucose or ^13^C5-glutamine in high iron conditions (0.625 mg/mL holotransferrin), n=4. **(c)** Relative abundance of α-KG mass isotopomers from ^13^C-glucose tracing, n=4. Schematic of tracing into α-KG from **(d)** ^13^C5-glutamine and **(e)** ^13^C6-glucose. Blue circles indicate ^13^C labelled atoms from ^13^C5-glutamine. Pink circles indicate ^13^C labelled atoms from ^13^C6-glucose. **(f)** NAD+/NADH ratio, n=4. Data are mean ± SEM. Statistics are: **(a, f)** matched two-tailed t-test; **(b)** two-way ANOVA with sample matching between metabolites but not between the two carbon tracers. **(c)** matched two-way ANOVAs with the Geisser-Greenhouse correction and the Sidak correction for multiple comparisons. *p < 0.05, **p < 0.01, ***p < 0.001, ****p < 0.0001.

**Supplementary Fig. 4.**
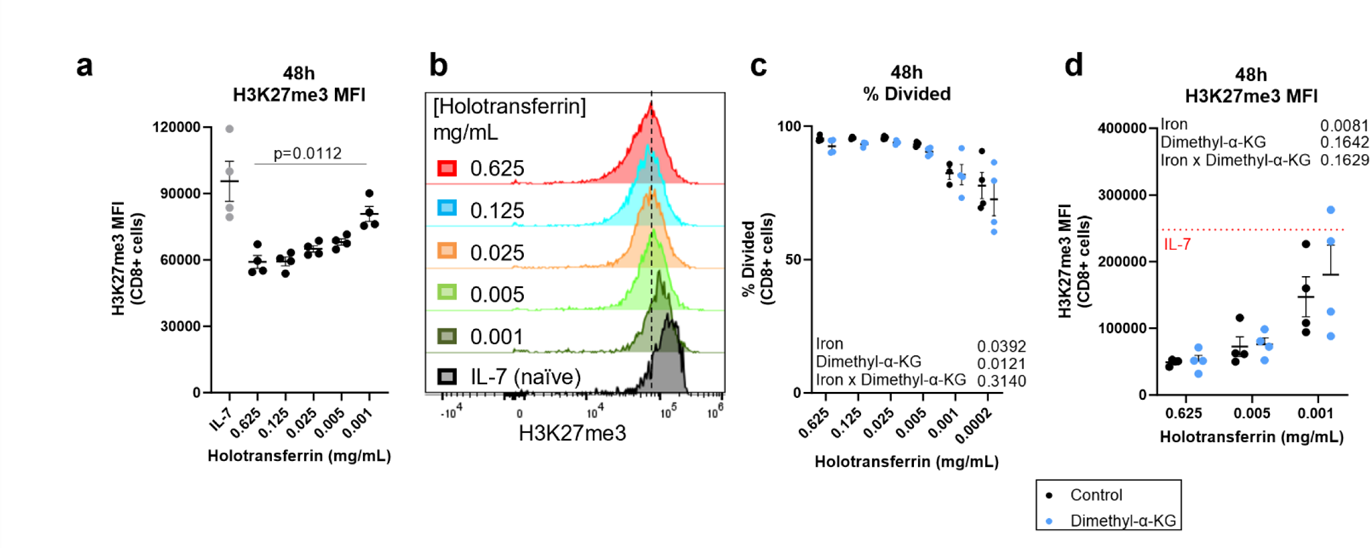
H3K27me3 accumulates in iron deprived CD8+ T-cells by 48h. CD8+ T-cells were activated as described in Fig. 1a. **(a-b)** H3K27me3 MFI measured at 48h, n=4. “Naïve” controls cells were cultured in IL-7 (5 ng/mL). **(c)** Percentage divided cells and **(d)** H3K27me3 MFI of cells cultured with or without dimethyl-α-KG (1 mM), n=4. Data are mean ± SEM. Histograms are normalised to mode. Statistics are: **(a)** matched one-way ANOVA with the Geisser-Greenhouse correction; **(c)** matched mixed effect analysis with the Geisser-Greenhouse correction; **(d)** two-way ANOVA with the Geisser-Greenhouse correction.

**Supplementary Fig. 5.**
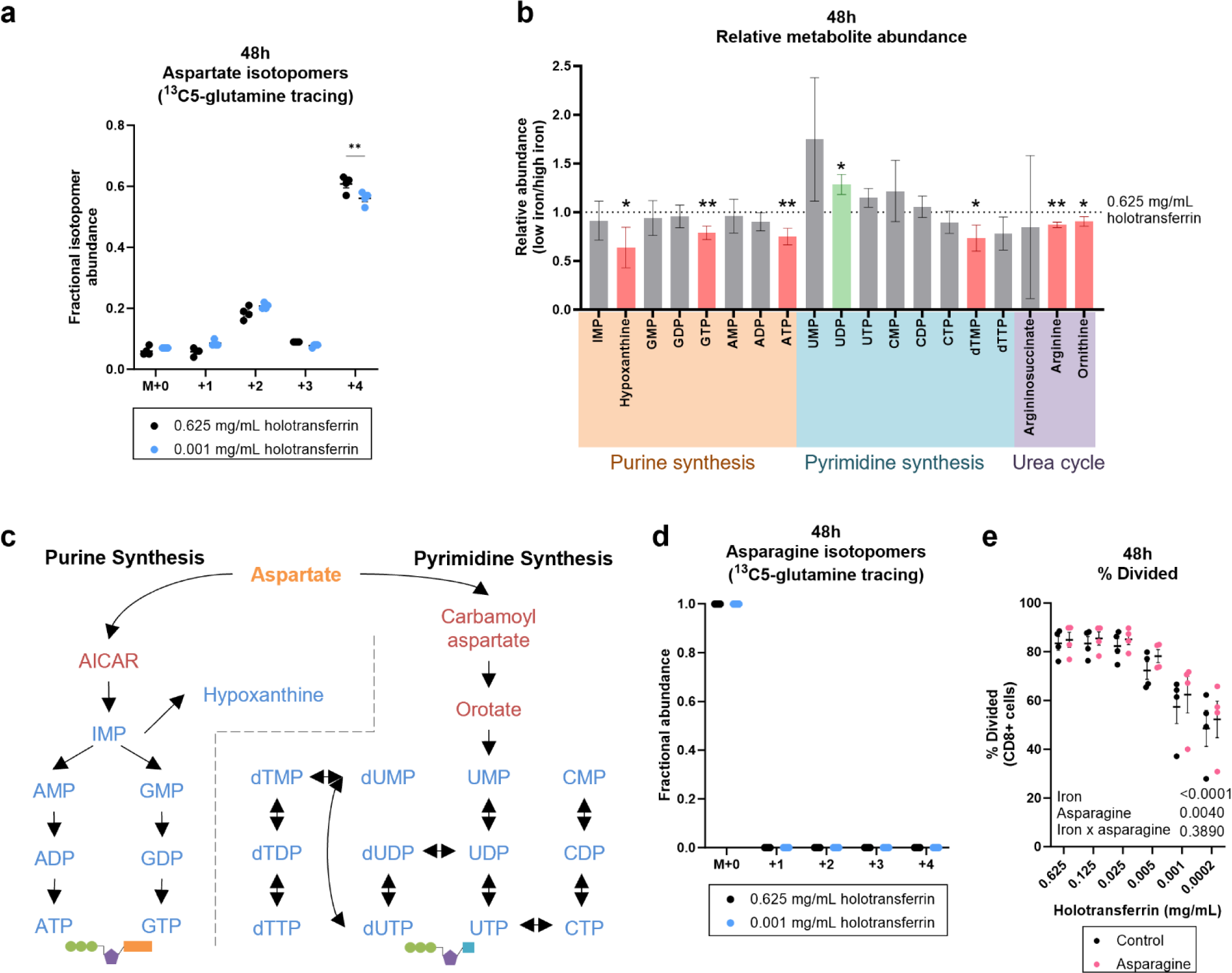
Iron deficient T-cells suppress aspartate utilising pathways. Isolated CD8+ T-cells were activated as described in Fig. 1a. For the ^13^C5-glutamine tracing experiments, T-cells were activated for 24h and then incubated in media containing ^13^C5-glutamine for a further 24h. **(a)** Aspartate and isotopomers labelled from ^13^C5-glutamine, n=4. M+1/2/3/4 indicate isotopomers derived from glutamine. **(b)** Relative metabolite abundance of T-cells in low iron (0.001 mg/mL holotransferrin) versus high iron (0.625 mg/mL holotransferrin) normalised to spiked in glutaric acid, n=4. **(c)** Aspartate is incorporated into ribonucleotides which can be interconverted between mono, di and tri-phosphorylated forms or converted to deoxy-ribonucleotides. **(d)** asparagine isotopomers labelled from ^13^C5-glutamine, n=4. M+1/2/3/4 indicate isotopomers derived from glutamine. **(e)** Division assessed with CTV with or without asparagine (100 µM), n=4. Data are mean ± SEM. Statistics are: **(a, d)** matched two-way ANOVAs with the Geisser-Greenhouse correction and the Sidak correction for multiple comparisons; **(b)** matched t-tests between 0.625 and 0.001 mg/mL holotransferrin conditions for each metabolite; **(e)** two-way ANOVA with the Geisser-Greenhouse correction.

**Supplementary Fig. 6.**
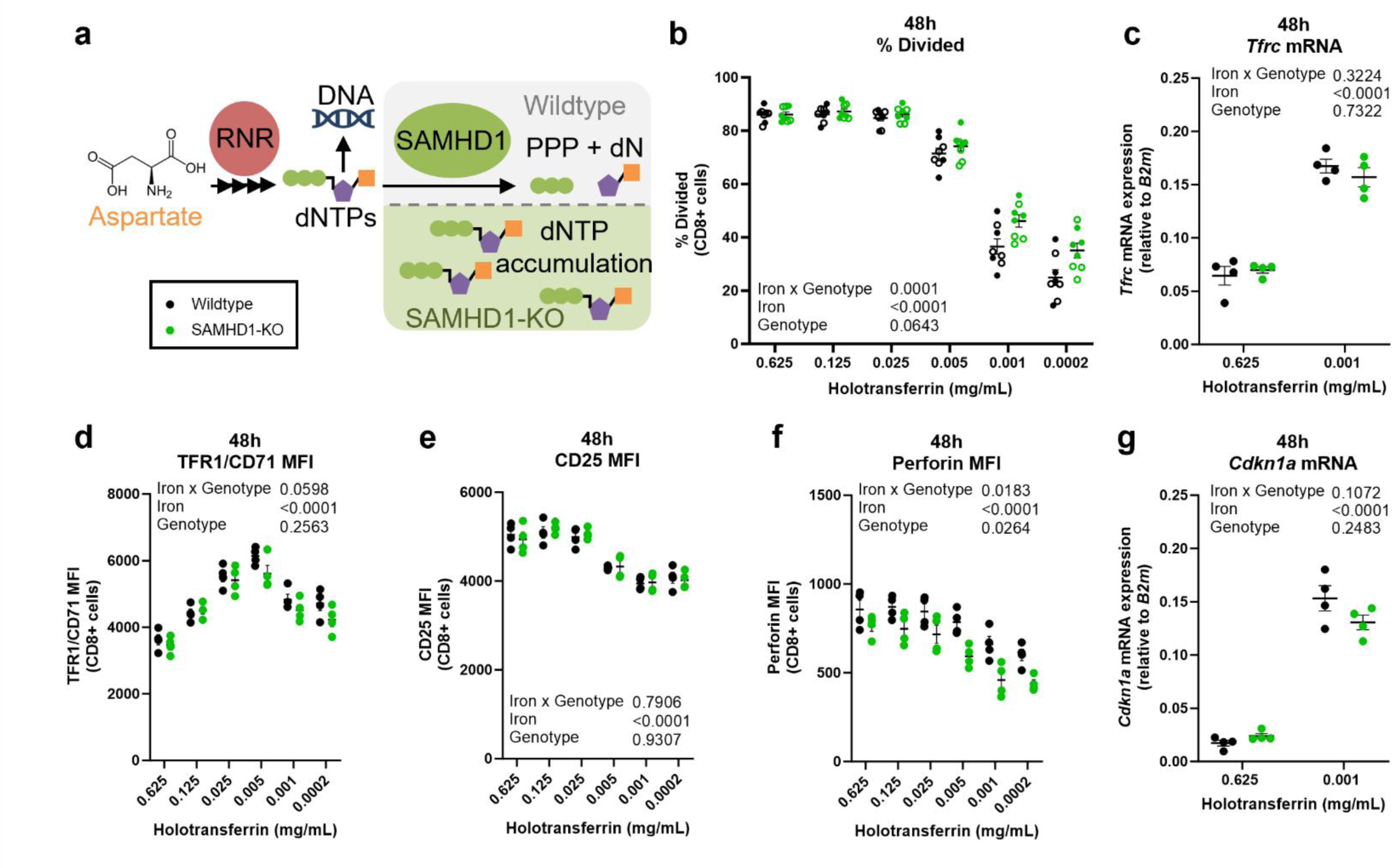
SAMHD1-KO T-cells show resistance to iron-deficiency suppressed proliferation. T-cells were isolated from SAMHD1-KO mice and wildtype littermate controls and were activated as described in Fig. 1a. **(a)** RNR enables dNTP production, SAMHD1 degrades dNTPs. SAMHD1-KO should result in dNTP accumulation. **(b)** Percentage divided cells measured using CTV. Data from independent experiments denoted by different symbols, n=8. *Tfrc*/TFR1/CD71 **(c)** mRNA and **(d)** surface protein MFI, n=4. **(e)** CD25 and **(f)** perforin MFI, n=4. **(g)** *Cdkn1a* mRNA expression by qPCR, n=4. Data are mean ± SEM. Statistics are: **(b-g)** sample matched two-way ANOVAs with the Geisser-Greenhouse correction applied for **(d-f)**.

**Supplementary Fig. 7.**
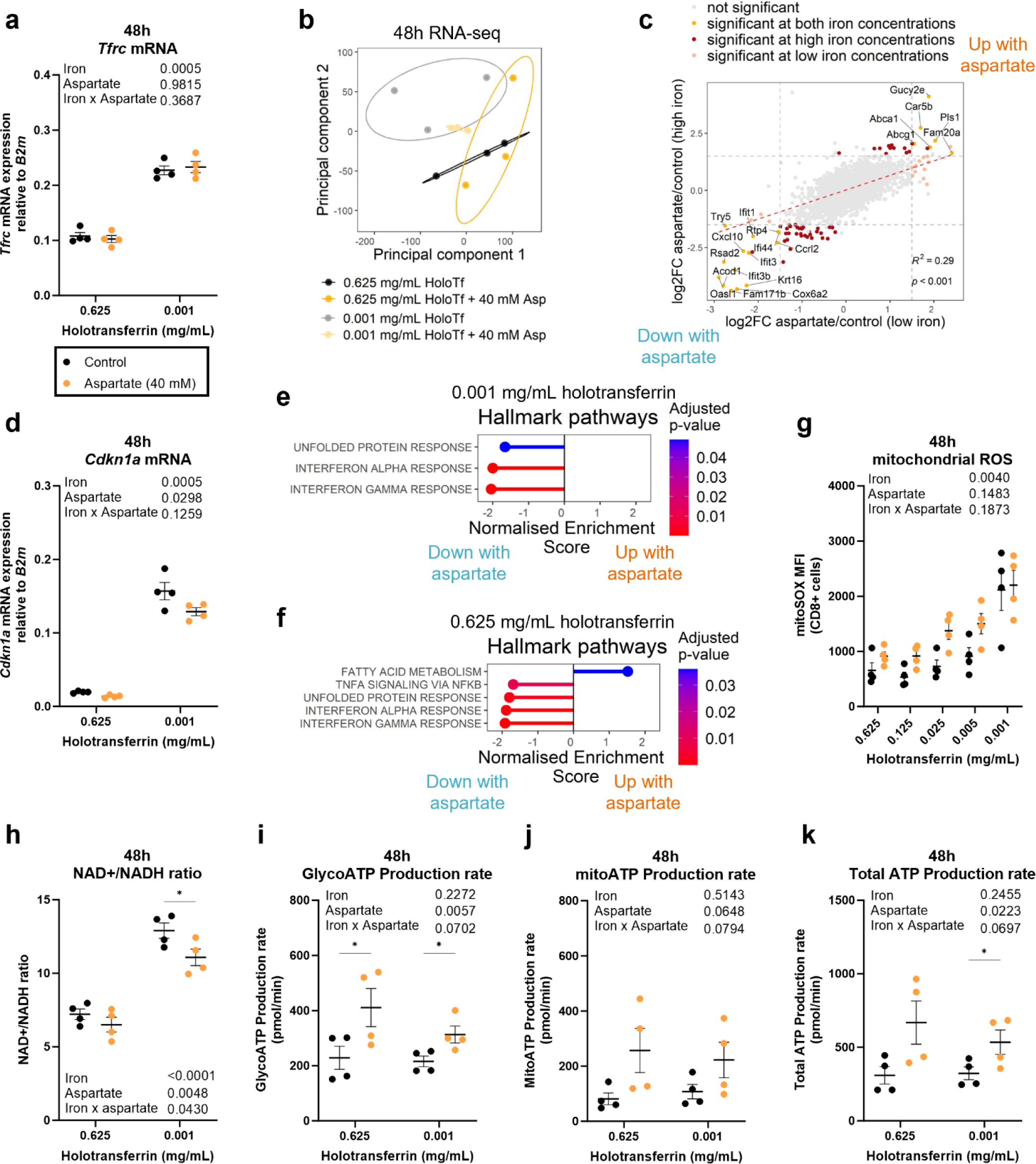
Aspartate has limited impact on cellular transcription but promotes a more metabolic state in CD8+ T-cells. CD8+ T-cells were activated as described in Fig. 1a with or without aspartate (40 mM). **(a)** *Tfrc* mRNA expression by qPCR, n=4. **(b)** RNA-seq PCA, n=4. **(c)** Correlation plot comparing the log_2_FC between aspartate and control conditions at either low iron or high iron conditions, n=4. Significance thresholds of |log_2_FC| > 1.5 and an FDR < 0.05 were applied. **(d)** *Cdkn1a* mRNA expression, n=4. GSEA of aspartate treated CD8+ T-cells versus control in **(e)** low iron conditions or **(f)** high iron conditions, n=4. **(g)** mROS MFI, n=4. **(h)** NAD+/NADH ratio, n=4. **(i)** Glycolytic, **(j)** mitochondrial and **(k)** total ATP production rate measured using the ATP rate seahorse kit, n=4. Data are mean ± SEM. Statistics are: **(a, d, g-k)** two-way ANOVAs with the Geisser-Greenhouse correction with Sidak’s test for multiple comparisons for **(h-k)**. **(c)** Pearson correlation R^2^ value. *p < 0.05.

## References

1 Ryan, D. G. et al. Coupling Krebs cycle metabolites to signalling in immunity and cancer. Nat Metab 1, 16–33 (2019). 10.1038/s42255-018-0014-7

2 Finley, L. W. S. What is cancer metabolism? Cell 186, 1670–1688 (2023). 10.1016/j.cell.2023.01.038

3 Prevalence, years lived with disability, and trends in anaemia burden by severity and cause, 1990-2021: findings from the Global Burden of Disease Study 2021. Lancet Haematol 10, e713-e734 (2023). 10.1016/s2352-3026(23)00160-6

4 Pasricha, S. R., Tye-Din, J., Muckenthaler, M. U. & Swinkels, D. W. Iron deficiency. Lancet 397, 233–248 (2021). 10.1016/s0140-6736(20)32594-0

5 Frost, J. N. et al. Hepcidin-Mediated Hypoferremia Disrupts Immune Responses to Vaccination and Infection. Med (N Y) 2, 164–179.e112 (2021). 10.1016/j.medj.2020.10.004

6 Andreini, C., Putignano, V., Rosato, A. & Banci, L. The human iron-proteome. Metallomics 10, 1223–1231 (2018). 10.1039/c8mt00146d

7 Dautry-Varsat, A. Receptor-mediated endocytosis: the intracellular journey of transferrin and its receptor. Biochimie 68, 375–381 (1986). 10.1016/s0300-9084(86)80004-9

8 Armitage, A. E. et al. Rapid growth is a dominant predictor of hepcidin suppression and declining ferritin in Gambian infants. Haematologica 104, 1542–1553 (2019). 10.3324/haematol.2018.210146

9 Shah, A., Frost, J. N., Aaron, L., Donovan, K. & Drakesmith, H. Systemic hypoferremia and severity of hypoxemic respiratory failure in COVID-19. Crit Care 24, 320 (2020). 10.1186/s13054-020-03051-w

10 Jabara, H. H. et al. A missense mutation in TFRC, encoding transferrin receptor 1, causes combined immunodeficiency. Nature Genetics 48, 74–78 (2016). 10.1038/ng.3465

11 Reina-Campos, M., Scharping, N. E. & Goldrath, A. W. CD8(+) T cell metabolism in infection and cancer. Nat Rev Immunol 21, 718–738 (2021). 10.1038/s41577-021-00537-8

12 Teh, M. R., Frost, J. N., Armitage, A. E. & Drakesmith, H. Analysis of Iron and Iron-Interacting Protein Dynamics During T-Cell Activation. Front Immunol 12, 714613 (2021). 10.3389/fimmu.2021.714613

13 Howden, A. J. M. et al. Quantitative analysis of T cell proteomes and environmental sensors during T cell differentiation. Nat Immunol 20, 1542–1554 (2019). 10.1038/s41590-019-0495-x

14 Wilkinson, N. & Pantopoulos, K. The IRP/IRE system in vivo: insights from mouse models. Front Pharmacol 5, 176 (2014). 10.3389/fphar.2014.00176

15 Levine, A. J., Hu, W. & Feng, Z. The P53 pathway: what questions remain to be explored? Cell Death Differ 13, 1027–1036 (2006). 10.1038/sj.cdd.4401910

16 Pereira, M. et al. Acute Iron Deprivation Reprograms Human Macrophage Metabolism and Reduces Inflammation In Vivo. Cell Rep 28, 498–511.e495 (2019). 10.1016/j.celrep.2019.06.039

17 Lai, Y. et al. Iron controls T helper cell pathogenicity by promoting glucose metabolism in autoimmune myopathy. Clin Transl Med 12, e999 (2022). 10.1002/ctm2.999

18 Kim, B. M., Choi, J. Y., Kim, Y. J., Woo, H. D. & Chung, H. W. Desferrioxamine (DFX) has genotoxic effects on cultured human lymphocytes and induces the p53-mediated damage response. Toxicology 229, 226–235 (2007). 10.1016/j.tox.2006.10.022

19 An, W. G. et al. Stabilization of wild-type p53 by hypoxia-inducible factor 1alpha. Nature 392, 405–408 (1998). 10.1038/32925

20 Ross, S. H., Rollings, C. M. & Cantrell, D. A. Quantitative Analyses Reveal How Hypoxia Reconfigures the Proteome of Primary Cytotoxic T Lymphocytes. Front Immunol 12, 712402 (2021). 10.3389/fimmu.2021.712402

21 Rath, S. et al. MitoCarta3.0: an updated mitochondrial proteome now with sub-organelle localization and pathway annotations. Nucleic Acids Res 49, D1541–d1547 (2021). 10.1093/nar/gkaa1011

22 Paul, B. T., Manz, D. H., Torti, F. M. & Torti, S. V. Mitochondria and Iron: current questions. Expert Rev Hematol 10, 65–79 (2017). 10.1080/17474086.2016.1268047

23 Turrens, J. F. Mitochondrial formation of reactive oxygen species. J Physiol 552, 335–344 (2003). 10.1113/jphysiol.2003.049478

24 Pearce, E. L., Poffenberger, M. C., Chang, C. H. & Jones, R. G. Fueling immunity: insights into metabolism and lymphocyte function. Science 342, 1242454 (2013). 10.1126/science.1242454

25 Bera, S. et al. Allosteric regulation of glutamate dehydrogenase deamination activity. Sci Rep 10, 16523 (2020). 10.1038/s41598-020-73743-4

26 Islam, M. S., Leissing, T. M., Chowdhury, R., Hopkinson, R. J. & Schofield, C. J. 2-Oxoglutarate-Dependent Oxygenases. Annu Rev Biochem 87, 585–620 (2018). 10.1146/annurev-biochem-061516-044724

27 Jaccard, A. et al. Reductive carboxylation epigenetically instructs T cell differentiation. Nature 621, 849–856 (2023). 10.1038/s41586-023-06546-y

28 Loenarz, C. & Schofield, C. J. Physiological and biochemical aspects of hydroxylations and demethylations catalyzed by human 2-oxoglutarate oxygenases. Trends Biochem Sci 36, 7–18 (2011). 10.1016/j.tibs.2010.07.002

29 Markolovic, S., Wilkins, S. E. & Schofield, C. J. Protein Hydroxylation Catalyzed by 2-Oxoglutarate-dependent Oxygenases. J Biol Chem 290, 20712–20722 (2015). 10.1074/jbc.R115.662627

30 Cribbs, A. P. et al. Histone H3K27me3 demethylases regulate human Th17 cell development and effector functions by impacting on metabolism. Proc Natl Acad Sci U S A 117, 6056–6066 (2020). 10.1073/pnas.1919893117

31 Russ, B. E. et al. Distinct epigenetic signatures delineate transcriptional programs during virus-specific CD8(+) T cell differentiation. Immunity 41, 853–865 (2014). 10.1016/j.immuni.2014.11.001

32 Villa, E., Ali, E. S., Sahu, U. & Ben-Sahra, I. Cancer Cells Tune the Signaling Pathways to Empower de Novo Synthesis of Nucleotides. Cancers (Basel) 11 (2019). 10.3390/cancers11050688

33 Morris, S. M., Jr. Regulation of enzymes of the urea cycle and arginine metabolism. Annu Rev Nutr 22, 87–105 (2002). 10.1146/annurev.nutr.22.110801.140547

34 Hope, H. C. et al. Coordination of asparagine uptake and asparagine synthetase expression modulates CD8+ T cell activation. JCI Insight 6 (2021). 10.1172/jci.insight.137761

35 Birsoy, K. et al. An Essential Role of the Mitochondrial Electron Transport Chain in Cell Proliferation Is to Enable Aspartate Synthesis. Cell 162, 540–551 (2015). 10.1016/j.cell.2015.07.016

36 Sullivan, L. B. et al. Supporting Aspartate Biosynthesis Is an Essential Function of Respiration in Proliferating Cells. Cell 162, 552–563 (2015). 10.1016/j.cell.2015.07.017

37 Schwartz, A. J. et al. Hepcidin sequesters iron to sustain nucleotide metabolism and mitochondrial function in colorectal cancer epithelial cells. Nature Metabolism 3, 969–982 (2021). 10.1038/s42255-021-00406-7

38 Pedley, A. M. & Benkovic, S. J. A New View into the Regulation of Purine Metabolism: The Purinosome. Trends Biochem Sci 42, 141–154 (2017). 10.1016/j.tibs.2016.09.009

39 Leone, R. D. et al. Glutamine blockade induces divergent metabolic programs to overcome tumor immune evasion. Science 366, 1013–1021 (2019). 10.1126/science.aav2588

40 Elia, I. et al. Tumor cells dictate anti-tumor immune responses by altering pyruvate utilization and succinate signaling in CD8(+) T cells. Cell Metab 34, 1137–1150.e1136 (2022). 10.1016/j.cmet.2022.06.008

41 Coggins, S. A., Mahboubi, B., Schinazi, R. F. & Kim, B. SAMHD1 Functions and Human Diseases. Viruses 12 (2020). 10.3390/v12040382

42 Franzolin, E. et al. The deoxynucleotide triphosphohydrolase SAMHD1 is a major regulator of DNA precursor pools in mammalian cells. Proc Natl Acad Sci U S A 110, 14272–14277 (2013). 10.1073/pnas.1312033110

43 Rehwinkel, J. et al. SAMHD1-dependent retroviral control and escape in mice. Embo j 32, 2454–2462 (2013). 10.1038/emboj.2013.163

44 Ruprecht, J. J. & Kunji, E. R. S. The SLC25 Mitochondrial Carrier Family: Structure and Mechanism. Trends Biochem Sci 45, 244–258 (2020). 10.1016/j.tibs.2019.11.001

45 Talpaz, M., Mercer, J. & Hehlmann, R. The interferon-alpha revival in CML. Ann Hematol 94 **Suppl 2**, S195–207 (2015). 10.1007/s00277-015-2326-y

46 Li, L. et al. Iron deprivation restrains the differentiation and pathogenicity of T helper 17 cell. J Leukoc Biol 110, 1057–1067 (2021). 10.1002/jlb.3ma0821-015r

47 Voss, K. et al. Elevated transferrin receptor impairs T cell metabolism and function in systemic lupus erythematosus. Sci Immunol 8, eabq0178 (2023). 10.1126/sciimmunol.abq0178

48 Wang, Z. et al. Iron Drives T Helper Cell Pathogenicity by Promoting RNA-Binding Protein PCBP1-Mediated Proinflammatory Cytokine Production. Immunity 49, 80–92.e87 (2018). 10.1016/j.immuni.2018.05.008

49 Henderson, S. A., Dallman, P. R. & Brooks, G. A. Glucose turnover and oxidation are increased in the iron-deficient anemic rat. Am J Physiol 250, E414–421 (1986). 10.1152/ajpendo.1986.250.4.E414

50 Shapiro, J. S. et al. Iron drives anabolic metabolism through active histone demethylation and mTORC1. Nat Cell Biol 25, 1478–1494 (2023). 10.1038/s41556-023-01225-6

51 Henning, A. N., Roychoudhuri, R. & Restifo, N. P. Epigenetic control of CD8(+) T cell differentiation. Nat Rev Immunol 18, 340–356 (2018). 10.1038/nri.2017.146

52 McNeill, L. A. et al. Hypoxia-inducible factor prolyl hydroxylase 2 has a high affinity for ferrous iron and 2-oxoglutarate. Mol Biosyst 1, 321–324 (2005). 10.1039/b511249b

53 Warburg, O. IRON, THE OXYGEN-CARRIER OF RESPIRATION-FERMENT. Science 61, 575–582 (1925). 10.1126/science.61.1588.575

54 Firkin, F. A. R., B. Interpretation of biochemical tests for iron deficiency: diagnostic difficulties related to limitations of individual tests. Aust Prescr 20, 74–76 (1997). 10.18773/austprescr.1997.063

55 Kelly, V., Al-Rawi, A., Lewis, D., Kustatscher, G. & Ly, T. Low Cell Number Proteomic Analysis Using In-Cell Protease Digests Reveals a Robust Signature for Cell Cycle State Classification. Mol Cell Proteomics 21, 100169 (2022). 10.1016/j.mcpro.2021.100169

56 Reyes, L. et al. -------A type I IFN, prothrombotic hyperinflammatory neutrophil signature is distinct for COVID-19 ARDS. Wellcome Open Res 6, 38 (2021). 10.12688/wellcomeopenres.16584.2

57 Muntel, J. et al. Surpassing 10 000 identified and quantified proteins in a single run by optimizing current LC-MS instrumentation and data analysis strategy. Mol Omics 15, 348–360 (2019). 10.1039/c9mo00082h

58 Wiśniewski, J. R., Hein, M. Y., Cox, J. & Mann, M. A “proteomic ruler” for protein copy number and concentration estimation without spike-in standards. Mol Cell Proteomics 13, 3497–3506 (2014). 10.1074/mcp.M113.037309

59 Perez-Riverol, Y. et al. The PRIDE database resources in 2022: a hub for mass spectrometry-based proteomics evidences. Nucleic Acids Res 50, D543–d552 (2022). 10.1093/nar/gkab1038

